# A unified model of short- and long-term plasticity: Effects on network connectivity and information capacity

**DOI:** 10.1101/2025.11.07.687160

**Authors:** Iiro Ahokainen, Marja-Leena Linne

## Abstract

Activity-dependent synaptic plasticity is a fundamental learning mechanism that shapes the connectivity and activity of neural circuits. Existing computational models of Spike-Timing-Dependent Plasticity (STDP) capture long-term synaptic changes with varying degrees of biological detail. A common approach is to neglect the influence of short-term dynamics on long-term plasticity, which may be an oversimplification for certain neuron types. Thus, there is a need for new models to investigate how short-term dynamics influence long-term plasticity. To address this gap, we introduce a novel phenomenological model, the Short-Long-Term STDP (SL-STDP) rule, which directly integrates the Tsodyks-Markram model of short-term dynamics with postsynaptic long-term plasticity. We fit the new model to recordings from layer 5 of the visual cortex and study how short-term plasticity affects the firing rate frequency dependence of long-term plasticity in a single synapse. Our analysis revealed that the pre- and postsynaptic frequency dependence of long-term plasticity plays a crucial role in shaping the self-organization of recurrent neural networks (RNNs) and their information processing through the emergence of sinks and source nodes. We applied the SL-STDP rule to RNNs and found that neurons in the SL-STDP network self-organize into distinct firing rate clusters, stabilizing the dynamics. We extended the experiments by including homeostatic balancing, namely weight normalization and excitatory-to-inhibitory plasticity, and observed differences in degree correlations between the SL-STDP network and a network without direct coupling between short-term and long-term plasticity. Finally, we evaluated how the modified connectivity affects the networks’ information capacity in reservoir computing tasks. The SL-STDP rule outperformed the uncoupled system in the majority of tasks, and including excitatory-to-inhibitory facilitating synapses further improved information capacity. Our study demonstrates that short-term dynamics–induced changes in the frequency dependence of long-term plasticity play a pivotal role in shaping network dynamics and link synaptic mechanisms to information processing in RNNs.

**Author summary:** The brain is a complex organ capable of developing, adapting and learning throughout life. Learning and development of the brain is facilitated by several different plasticity mechanisms that act on different brain areas and timescales. Computational modeling of these plasticity mechanisms not only help us to understand the working principles of the brain but can provide us useful algorithms for future computing devices. In this work, we develop a new activity-dependent plasticity model that acts locally in a synapse and combines two timescales of plasticity into one set of equations. We investigate the effects of this new synapse model on recurrently connected neural networks and observe changes in neural activity and connectivity compared to traditional approaches. We further evaluate the model by performing information capacity tests that are related to the working memory of the circuit. We find improved information capacity, indicating enhanced computational performance of the proposed model. Our study emphasizes how combining the two timescales of plasticity may be important for the development of neural circuits and working memory.

## Introduction

Learning and memory in biological neuronal networks depend on activity-dependent changes in synaptic efficacy and on the resulting reorganization of network connectivity. Long-term potentiation (LTP) and long-term depression (LTD) provide two central mechanisms for such reorganization by strengthening and weakening synapses over long periods of time. A widely used phenomenological principle describing these long-term synaptic modifications is spike-timing-dependent plasticity (STDP), in which correlations between pre- and postsynaptic spiking determine long-term changes in synaptic efficacy [1, 2]. Together with homeostatic mechanisms, LTP and LTD can form stable assembly formations [3, 4], suggesting that long-term plasticity can drive recurrent neural networks (RNNs) in the brain toward connectivity profiles that support robust information processing. Importantly, synaptic transmission is also shaped on shorter timescales through mechanisms such as short-term depression and facilitation. These short-term synaptic dynamics can modulate neuronal communication and influence the induction of long-term plasticity. In particular, recurrent network models using STDP, short-term depression, and homeostatic plasticity have demonstrated that the coexistence of these mechanisms can critically affect the formation, retention, and reorganization of cell assemblies (see e.g. [4, 5]).

Phenomenological models of STDP commonly describe long-term synaptic modifications using temporal kernels or synaptic traces that capture correlations between pre- and postsynaptic spiking activity [6–9]. Postsynaptic traces or kernels are typically associated with postsynaptic calcium dynamics [10, 11], whereas explanations for presynaptic traces are less consistent. Possible interpretations include presynaptic calcium concentrations, glutamate levels in a synaptic cleft, or the fraction of open NMDA receptors [8, 9, 12], each of which may be modulated by presynaptic short-term dynamics. Together, these observations suggest that presynaptic short-term dynamics may directly influence long-term synaptic plasticity. Supporting this hypothesis, Froemke and colleagues showed that short-term depression directly affects long-term synaptic modifications in layer 2/3 synapses of visual cortical slices [13–15].

Despite these findings, only a limited number of STDP models explicitly incorporate short-term synaptic dynamics into long-term plasticity mechanisms. Motivated by the suppression models of Froemke and colleagues [13, 14], a few STDP variants incorporating suppressive short-term depression have been proposed [12, 16–19], whereas short-term facilitation has primarily been explored in calcium-based STDP models [20–22]. The application of these plasticity models in network-level simulations has been limited. As an exception, a recent study [23] integrated a biophysically detailed calcium-based plasticity model [22] into a large-scale cortical network model to investigate the role of plasticity at the microcircuit level. Additionally, Costa and colleagues developed a model of presynaptic long-term plasticity in which long-term modifications alter short-term dynamics, whereas short-term dynamics do not directly influence long-term plasticity induction [24]. Thus, the direction of interaction differs fundamentally from that considered in the present work. Taken together, the impact of models directly coupling short-term dynamics to long-term plasticity on connectivity and information processing in RNNs remains largely unexplored.

A central computational function associated with RNNs is working memory, defined as the ability of a system to temporarily retain information for complex input–output transformations [25]. Different hypotheses have been proposed regarding the mechanisms underlying working memory. The most common hypothesis assumes that working memory emerges from sustained spiking activity and attractor dynamics in recurrent cortical circuits [26]. Some approaches suggest that other biophysical mechanisms, such as short-term dynamics [27, 28] or intrinsic adaptive currents [29], could act directly as the locus of working memory. In this work, we focus on working memory and information capacity arising from sustained spiking activity, and investigate how synaptic self-organization driven by directly coupled short- and long-term plasticity shapes information capacity in RNNs.

To further study interactions between short- and long-term plasticity at both the single synapse and network levels, we develop a novel STDP model that directly combines the Tsodyks-Markram (TM) model of short-term dynamics [27, 30] with the Triplet STDP model [8]. This simplified approach enables us to compare the effects of short-term dynamics to long-term plasticity in a controlled manner. We refer to this new model as Short-Long-Term Spike-Timing-Dependent Plasticity (SL-STDP) rule, as it incorporates short-term depression and facilitation together with LTP and LTD within a unified set of equations. We start by fitting the SL-STDP model to layer 5 pyramidal neuron data from the rat visual cortex [31], and then compare the expected rate of change of the synaptic weight under stationary and uncorrelated Poisson spike statistics with that observed for the minimal all-to-all Triplet model [8]. Comparing the expected rates of change of the synaptic weights helps us to understand how the interaction between short-term dynamics and long-term plasticity shapes the firing rate frequency dependence of long-term plasticity. Next, we construct two RNNs [32], one incorporating the SL-STDP synapse and the other using uncoupled TM and Triplet synapses. We then compare their in- and out-degree distributions and network activity with different homeostatic balancing methods [33], namely weight normalization [34–36] and excitatory-to-inhibitory plasticity [37–41]. Different homeostatic balancing methods are commonly applied in RNNs, and understanding how the new plasticity model interacts with them is therefore critical. Finally, we evaluate networks’ information capacity [42] in reservoir computing tasks [43, 44]. We first allow the networks to self-organize with the plasticity rules, after which we evaluate information capacity on increasingly challenging firing rate encoded fading memory and nonlinear transformation tasks [42, 45].

We found that the SL-STDP synapse exhibits versatile pre- and postsynaptic firing rate frequency dependencies that vary depending on the balance between short-term depression and facilitation. We argue that these frequency dependencies have a significant influence on the self-organization of RNNs through the formation of sink and source nodes. Importantly, we found that directly coupling short-term dynamics to long-term plasticity is sufficient alone to prevent part of the positive feedback loops that otherwise lead to runaway synaptic potentiation of all excitatory-to-excitatory synapses. Instead, the network self-organizes into distinct firing rate clusters. Further analyses incorporating synaptic weight normalization and excitatory-to-inhibitory plasticity revealed differences in out-degree and in-degree correlations between the SL-STDP and the uncoupled TM-Triplet model. Information capacity analyses further demonstrated that the SL-STDP model outperforms the uncoupled TM-Triplet model across multiple fading memory and nonlinear transformation tasks. In particular, the SL-STDP network model with a small fraction of facilitating excitatory-to-inhibitory connections achieved the highest overall information capacity values.

## Results

### Unifying short-term plasticity and long-term plasticity through a shared state

We propose a new synapse model by directly combining short-term plasticity with long-term plasticity through a shared latent variable *y*(*t*) in order to study the direct impact of short-term dynamics on long-term plasticity. Throughout the rest of this article, we refer to this new plasticity model as the Short-Long-Term

Spike-Timing-Dependent Plasticity (SL-STDP) rule. The state variable *y*(*t*) is controlled by the presynaptically released neurotransmitter and is usually referred to as effective or active resources [30, 46]. The presynaptic part of the synapse model then follows the typical Tsodyks-Markram dynamics of vesicle depletion *x*(*t*) and release probability *u*(*t*) [27, 30]:

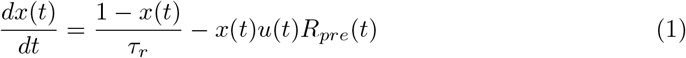

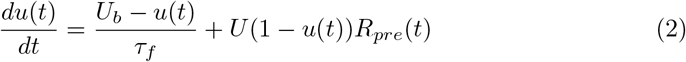

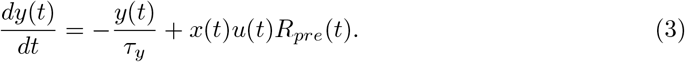

At each presynaptic spike *t*_*pre*_, the amount of readily releasable neurotransmitter *x*(*t*) is reduced by *x*(*t*)*u*(*t*), and the release probability *u*(*t*) is increased by the maximum of *U* from the baseline release probability *U*_*b*_. The presynaptic spike train is defined with a neural response function *R*_*pre*_(*t*) = Σ *δ*(*t* − *t*_*pre*_). The timescale of short-term presynaptic dynamics is determined by the time constants for facilitation, depression, and the synaptic response to the released neurotransmitter, denoted by *τ*_*f*_, *τ*_*r*_, and *τ*_*y*_, respectively. Whether the short-term dynamics are facilitating, depressing or mixed depends on the parameters *τ*_*f*_, *τ*_*r*_, *U* and *U*_*b*_ [30, 47–49]. Typical parameter sets for each case are shown in Table 1 [48].

**Table 1.** Example parameter sets for different short-term plasticity dynamics. [48].

| Short-term dynamics type | $\tau_r$ (ms) | $\tau_f$ (ms) | $U$ | $U_b$ |
| --- | --- | --- | --- | --- |
| Depression | 500 | 50 | 0.05 | 0.5 |
| Facilitation - Depression | 200 | 200 | 0.3 | 0.25 |
| Facilitation | 50 | 500 | 0.15 | 0.15 |

The postsynaptic states *z*_1_(*t*) and *z*_2_(*t*) evolve as

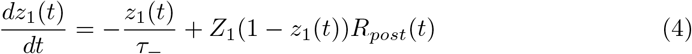

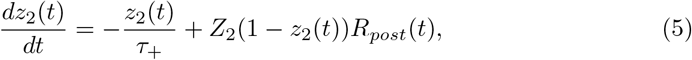

where *R*_*post*_(*t*) is the postsynaptic neural response function, and *τ*_−_ and *τ*_+_ are the time constants for the Long-Term Depression (LTD) and Long-Term Potentiation (LTP) states *z*_1_(*t*) and *z*_2_(*t*), respectively. We include gain control defined by *Z*_1_ and *Z*_2_ for the states *z*_1_(*t*) and *z*_2_(*t*), respectively. Gain control effectively limits the number of spikes that contribute to the integration of postsynaptic states. The inclusion of gain control on the postsynaptic side can be biologically motivated, e.g. by Ca^2+^-dependent inactivation [50–52], K^+^-mediated spike attenuation [14, 53, 54] or any other immediate regulatory mechanism.

Lastly, an LTD update Δ*w*^−^(*t*) and an LTP update Δ*w*^+^(*t*) are defined as

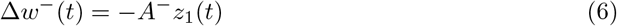

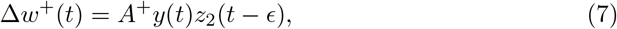

where *A*^−^ is the update scale for the pair-based LTD and *A*^+^ is the scale of update for the triplet-based LTP. A small *ϵ* ensures that the LTP update is indeed triplet-based [8]. Note that the LTP update directly depends on the state *y*(*t*), which is affected by the short-term plasticity dynamics in Eqs (1-3), which is not usually the case in many STDP learning systems. On each presynaptic spike, the current connection weight *w* is then updated as

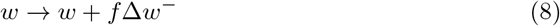

and on each postsynaptic spike as

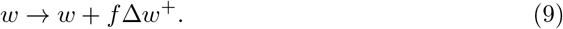

Here, *f* is the learning rate for both LTP and LTD updates. We also include a hard cap, *w*_max_, to limit the unbounded growth of *w*. We chose this particular form of updates in Eqs (6-7) because a pair-based LTD and a triplet-based LTP are enough to explain the majority of the frequency dependence of weight updates in the rat L5 visual cortex *ex vivo* [8]. However, alternative combinations of pair- or triplet-based updates could be used if needed. In addition, multiplicative or power law updates [7] could be included but were omitted for clarity.

To analyze how the connection weight *w* evolves depending on short-term dynamics, we derive the expected rate of change of the synaptic weight, which we refer to as the mean synaptic drift. The mean synaptic drift, 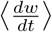, provides a qualitative explanation of the dynamics of *w* at a single synapse over longer timescales. Assuming two uncorrelated and independent Poisson spike trains, one presynaptic and one postsynaptic, with mean firing rates *λ*_*x*_ and *λ*_*z*_, respectively, one obtains the mean synaptic drift

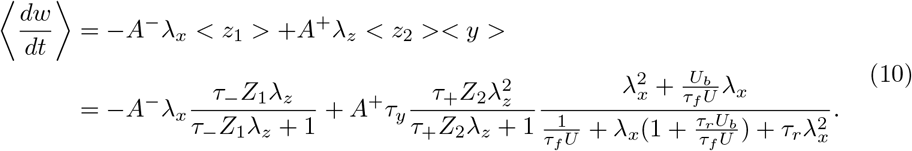

Here <·> denotes a steady-state value. From Eq (10), we observe that the positive update part (second term on the right) does not scale linearly with *λ*_*x*_, in contrast to other STDP rules that follow Bienenstock-Cooper-Munro (BCM) model [55] (see, e.g. [8, 9, 56]). Instead, there exists a maximum 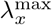, whose position depends on the steady state value < *y* >. Thus, depressing and facilitating synapses have different preferred firing rate regimes for maximal potentiation, as illustrated in Fig 1.

**Fig 1.**
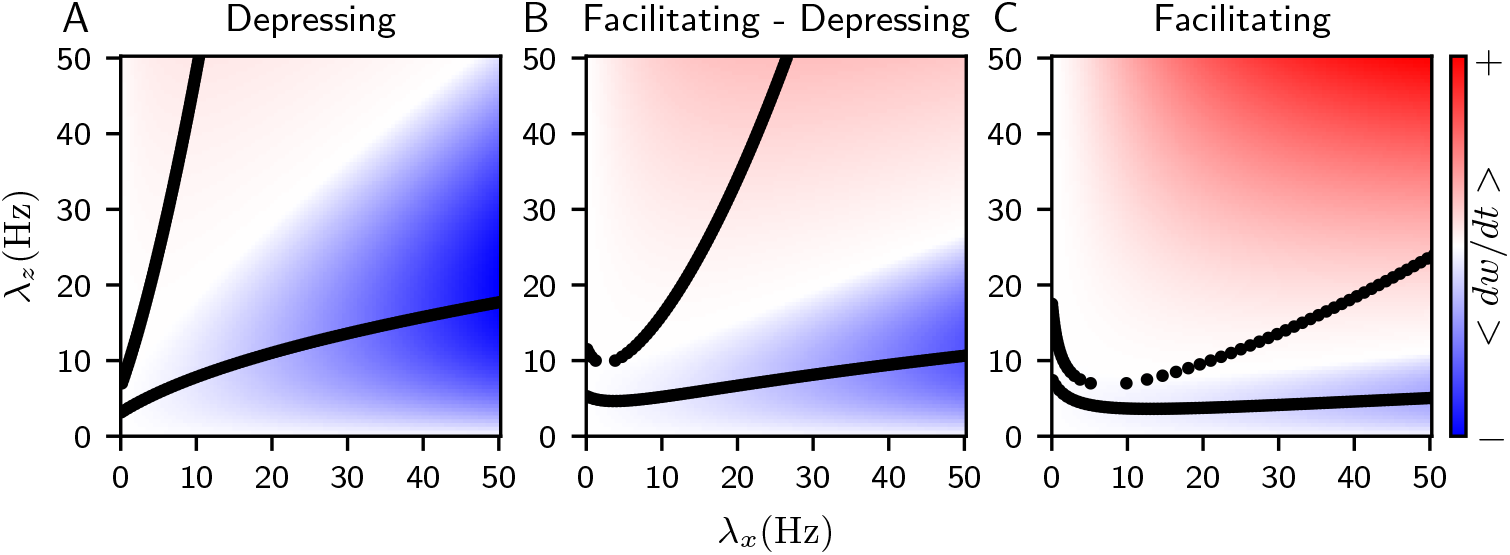
The SL-STDP synapse model exhibits varying frequency responses depending on short-term dynamic parameters. The mean drift of the connection weight *w* is shown as a function of the presynaptic firing rate *λ*_*x*_ and the postsynaptic firing rate *λ*_*z*_. Red indicates net LTP, blue indicates net LTD, and white denotes the regime where LTP and LTD are balanced. Dotted black lines represent the nullclines of the system, denoting local maxima and minima of Eq (10). (**A**) Short-term depression-dominated synapse exhibits strong LTD, while (**C**) facilitating synapse shows enhanced LTP. (**B**) Balanced facilitation and depression display features from both facilitating and depressing synapses. The short-term plasticity parameters for panels (**A**-**C**) are given in Table 1. The remaining parameters are identical across panels (**A**-**C**): *A*^−^ = 0.02, *A*^+^ = 1, *τ*_*y*_ = 16.8 ms, *τ*_−_ = 100 ms, *τ*_+_ = 50 ms, *Z*_1_ = 0.5 and *Z*_2_ = 0.3.

To further clarify the difference between depressing and facilitating synapses, consider the case where *λ*_*z*_ is large. For a depressing synapse with large *λ*_*x*_, < *y* > → 1*/τ*_*r*_, assuming *τ*_*f*_ → 0. Therefore, we obtain the relation 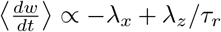, which implies that stronger depression (larger *τ*_*r*_) results in reduced potentiation, and < *y* > → *λ*_*x*_, and therefore 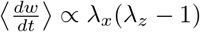. This means that when the postsynaptic when *λ*_*x*_ dominates over *λ*_*z*_, there is net LTD and vice versa. In contrast, for a facilitating synapse, we consider the limits *τ*_*r*_ → 0 and *τ*_*f*_ → ∞. In this case, firing rate is sufficiently small, the synapse exhibits more LTD than LTP, and the LTD zone scales linearly with *λ*_*x*_, similar to the Triplet rule (Eq (16)). This phenomenon is illustrated in Fig 1C. Also, the LTP zone scales linearly with both in *λ*_*x*_ and *λ*_*z*_, indicating that there is no local maximum for the potentiation in the facilitating synapse. In practice, with finite parameter values, the region of maximal potentiation shifts toward larger *λ*_*x*_ values when transitioning from a depression-dominated synapse to a facilitation-dominated synapse (Fig 1A-C). Note that Fig 1 shows 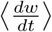 with a single set of postsynaptic parameters and relative amplitudes, and the LTP and LTD regions will change when e.g. *Z*_1_ and *Z*_2_ are modified.

### Fitted SL-STDP model exhibits balanced firing rate frequency dependencies

We fit the SL-STDP model to experimental pyramidal neuron data from the rat layer 5 visual cortex [31]. We utilize a random firing dataset in addition to the fixed Δ*t* = *±*10 ms pre-post spike interval to avoid overfitting to the Δ*t* = *±* 10 ms data set (see Methods). The model fits are shown in Fig 2, and the fitted parameters and the fitting error are reported in Table 2. For model fitting, we fix the short-term plasticity time constants to *τ*_*r*_ = 200 ms and *τ*_*f*_ = 50 ms, as depressing synapses are commonly observed between layer 5 pyramidal neurons [24, 48, 57, 58]. All other parameters in Eqs (1-7) are fitted. In particular, the fitting procedure resulted in a time constant *τ*_*y*_ = 16.54 ms for the state *y*(*t*). Note that the same presynaptic spike efficacy *y*(*t*) is used for the input current in Eq (39). The exact interpretation of *y*(*t*) is not at the focus since the model aims to explain functional consequences of the coupling of short-term and long-term plasticity rather than the exact biological mechanism.

**Table 2.** Fitted SL-STDP model parameters. Short-term plasticity time constants were fixed at *τ*_*r*_ = 200 ms and *τ*_*f*_ = 50 ms, and all other parameters were fitted. The last column on the right shows the total fitting error given by Eq (19). The average standard error of the mean for the experimental data points in Fig 2A is 14.6 %, whereas the fitted model exhibits a mean difference of 12.1 % between the model’s values and the experimental data points.

| $U$ | $U_b$ | $\tau_y$ (ms) | $\tau_-$ (ms) | $Z_1$ | $A^-$ | $\tau_+$ (ms) | $Z_2$ | $A^+$ | Error |
| --- | --- | --- | --- | --- | --- | --- | --- | --- | --- |
| 0.2029 | 0.7019 | 16.54 | 100.0 | 0.4102 | 0.0166 | 63.30 | 0.1158 | 0.8499 | 487.9 |

**Fig 2.**
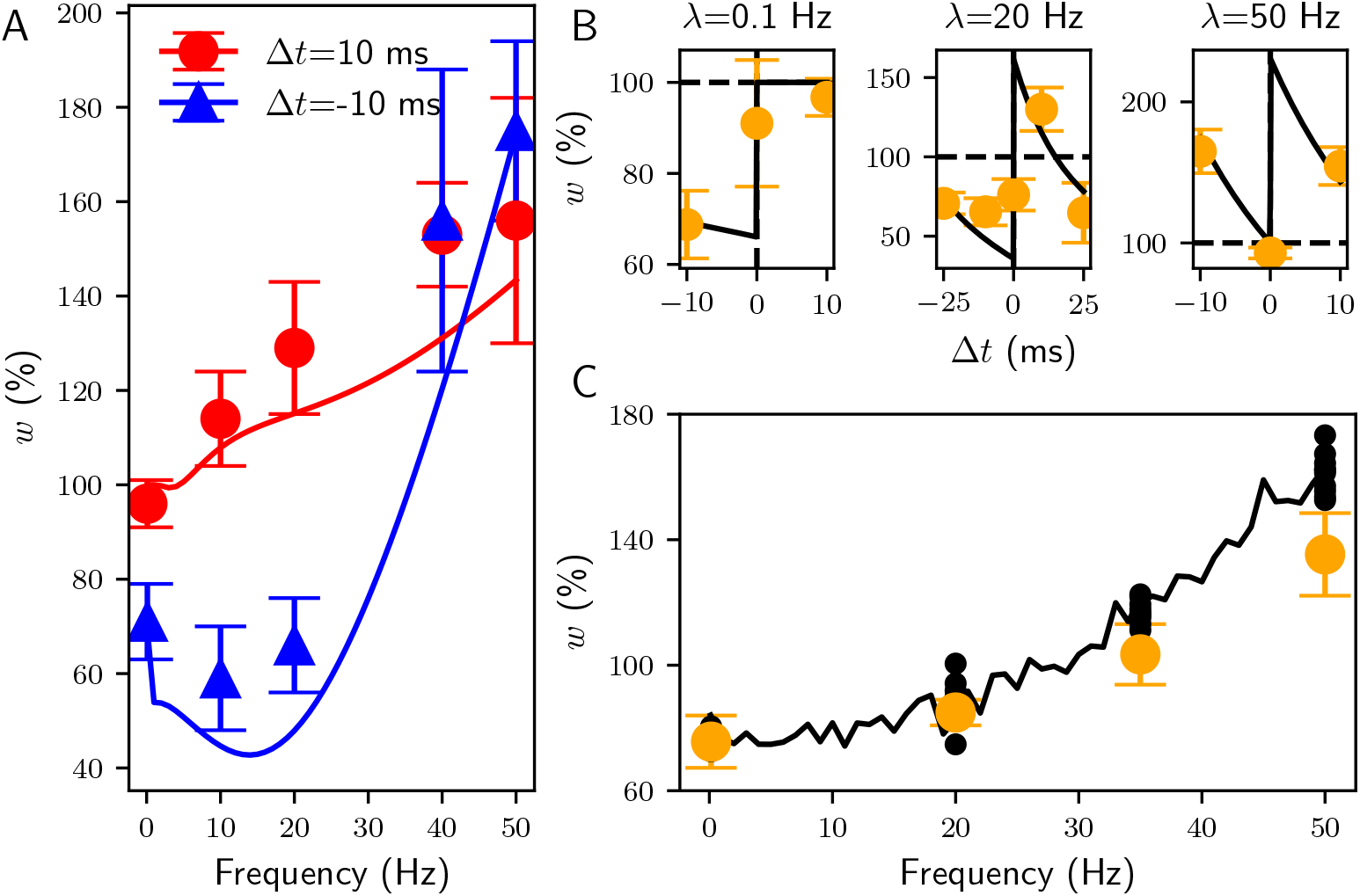
SL-STDP model fit to the long-term plasticity induction protocol [31]. (**A**) Change of *w* in response to varying interspike intervals resulting to the nonlinear frequency response with fixed Δ*t* = *±* 10 ms. Dots and triangles with the standard deviations is the data from [31] and lines are the fits. (**B**) The update window with varying Δ*t* given three different frequencies 0.1 Hz, 20 Hz, and 50 Hz, from left to right. (**C**) Model fit to the randomized Δ*t*. As in [31], Δ*t* was drawn from a Gaussian distribution with a mean zero and a standard deviation 7 ms. Orange error bars show the experimental data and black dots show all the realizations of model fits. The black line shows one example fit over the whole frequency range. See Table 2 for the fitted parameter values.

It is necessary to repeat the same fitting procedure for the minimal all-to-all Triplet model [8] for later comparison. The original Triplet model fitting did not include the stochastic data set nor all the data points shown in Fig 2B [8]. In addition, we closely followed the experimental long-term plasticity induction protocol presented in [31] in contrast to [8], since the relaxation time between 5-5 spike trains may be important for the short-term dynamics (compared to a scenario where all pre- and postsynaptic spikes are given with a fixed interval [8]). Thus, the newly fitted minimal all-to-all Triplet model is used as a basis for comparison throughout the remainder of the article. The model fits for the Triplet model are found in S1. Details of the fitting procedures for both models are described in the Methods section.

Fig 3 shows the example response of the SL-STDP synapse model when stimulated with presynaptic and postsynaptic Poisson distributed spike trains with rates *λ* = 50 Hz. As presynaptic traces (Fig 3B) and postsynaptic traces (Fig 3C) evolve in response to the Poisson spike trains (Fig 3D), the weight parameter *w* evolves slowly (Fig 3A). The synapse model with Table 2 parameter values results in a predominantly depressing synapse as can be inferred from the *y*(*t*) trace in Fig 3B. Note that the postsynaptic traces in Fig 3C do not increase unbounded due to the gain control Eq (4-5). Thus, the later spikes in a burst have less importance, which aligns well with the revised suppression model [14].

**Fig 3.**
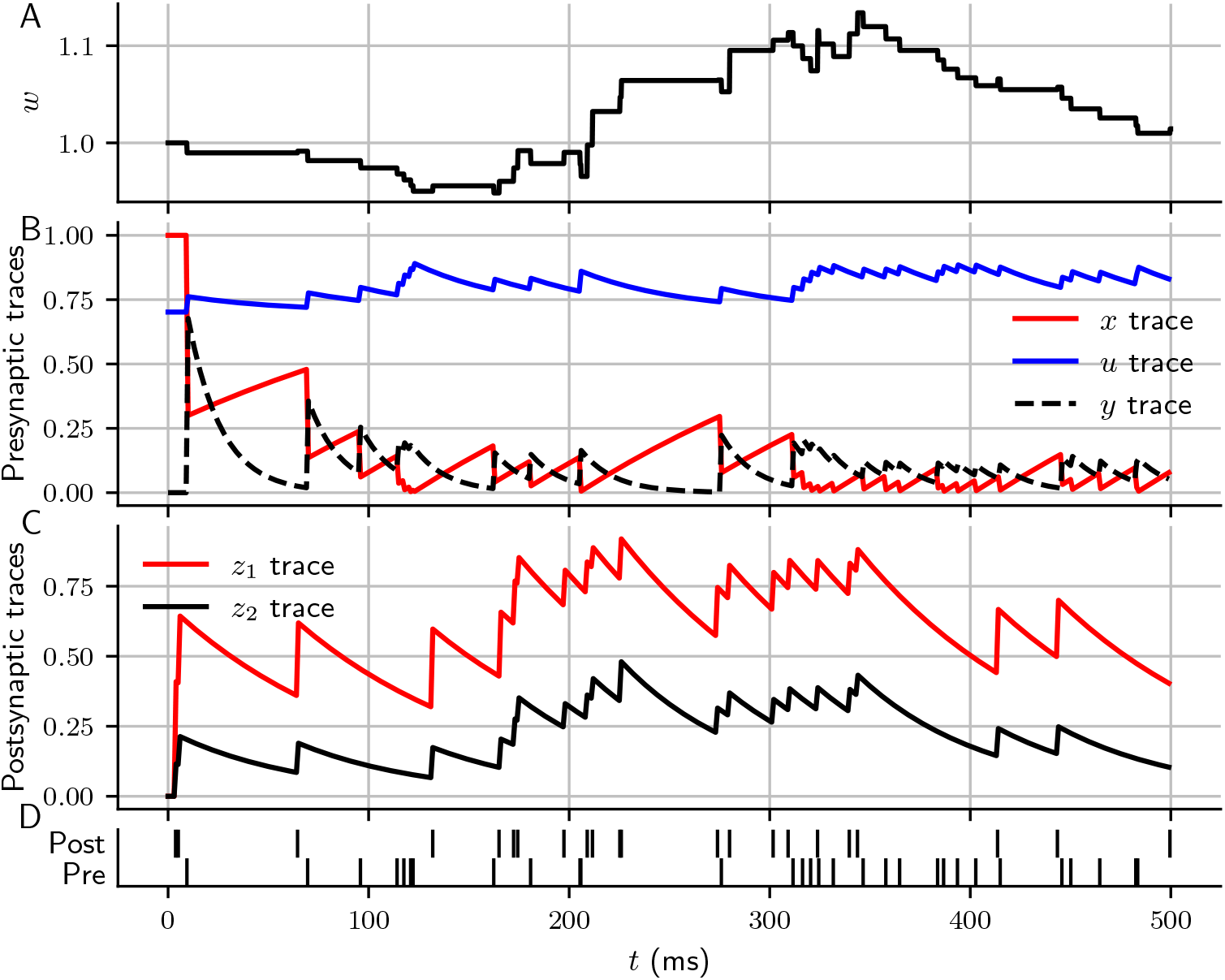
Example evolution of the SL-STDP synapse model. (**A**) The synaptic weight *w* evolves according to Eqs (8-9) as presynaptic traces (**B**) and postsynaptic traces (**C**) respond to presynaptic and postsynaptic spikes (**D**). Eqs (1-5) were solved in an event-based manner. The used parameters are shown in Table 2.

The mean drift of *w* (Eq (10)) for the fitted Triplet model and the SL-STDP model is shown in Fig 4. Surprisingly, the scale of 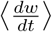 differs significantly between the two models, despite both being fitted to the same data using the same protocol. The maximal LTP in the Triplet model is two orders of magnitude greater than the maximal LTD (Fig 4A). In contrast, the SL-STDP model exhibits balanced amplitudes for LTP and LTD (Fig 4B). In addition, the shapes of the LTD and LTP zones differ substantially between the two models. In the Triplet model, higher presynaptic firing rates result in increasing LTP when *λ*_*z*_ exceeds a fixed threshold 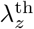, and in LTD when 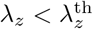. Note that 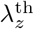 could be made adaptive following the BCM theory [8, 55]. The SL-STDP model does not necessarily lead to increased LTP even when *λ*_*z*_ is high and *λ*_*x*_ is increasing. This stands in a stark contrast to the Triplet model. For comparisons of other STDP models, see [59] for the mean drift matrix of the voltage-based rule by Clopath et al. [9] and the calcium-based rule by Graupner and Brunel [60].

**Fig 4.**
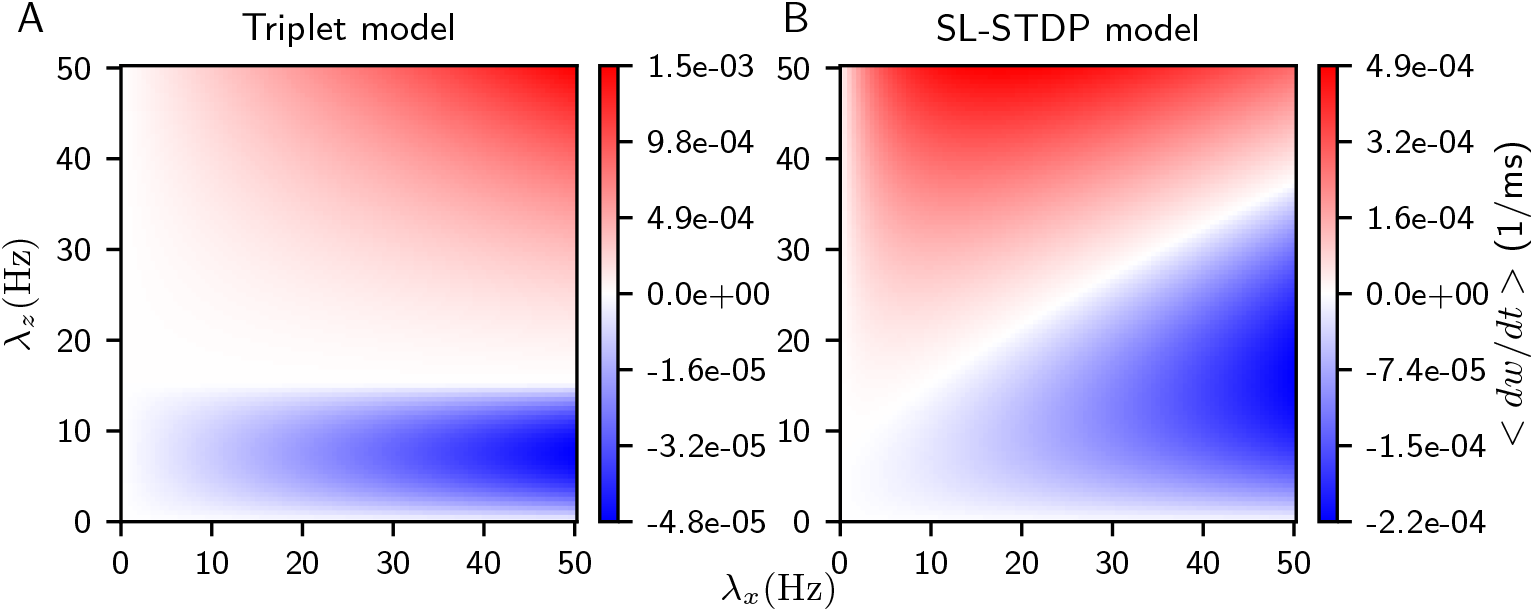
The mean drift 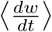 for the fitted models. Both models were first fitted to experimental data (see Methods), and 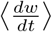 was then computed as a function the of presynaptic firing rate *λ*_*x*_ and the postsynaptic firing rate *λ*_*z*_. (**A**) 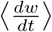 of the fitted Triplet model according to Eq (16). (**B**) 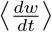 of the SL-STDP model according to Eq (10). Red areas indicate the net LTP regimes, and blue areas indicate the net LTD regimes.

### SL-STDP synapse and Triplet synapse generate different connectivity profiles in RNNs

We hypothesize that the balanced LTP and LTD zones, emerging from the direct coupling of short-term dynamics with long-term plasticity, promote richer network dynamics through the formation of source and sink nodes in recurrent neural networks (RNNs). A simplified illustration of this concept is shown in Fig 5. First, a small RNN in Fig 5A, consisting of four nodes, exhibits varying firing rates of strength I to III. Applying weight updates of form Fig 4B results in an RNN network shown in Fig 5B. Due to the symmetry of the weight update matrix, the relatively more quiescent nodes I and II morph into source nodes, while the stronger nodes III become sink nodes. In contrast, updates of the form shown in Fig 4A do not weaken the connections from nodes III to node II but instead strengthen them. Thus, the resulting RNN in Fig 5C contains only one clear source node I, while nodes II and III form a self-enforcing positive feedback loop. This simplified example that neglects input-output correlations and exact spike times demonstrates how the shape of the weight update matrix critically influences the connectivity in RNNs.

**Fig 5.**
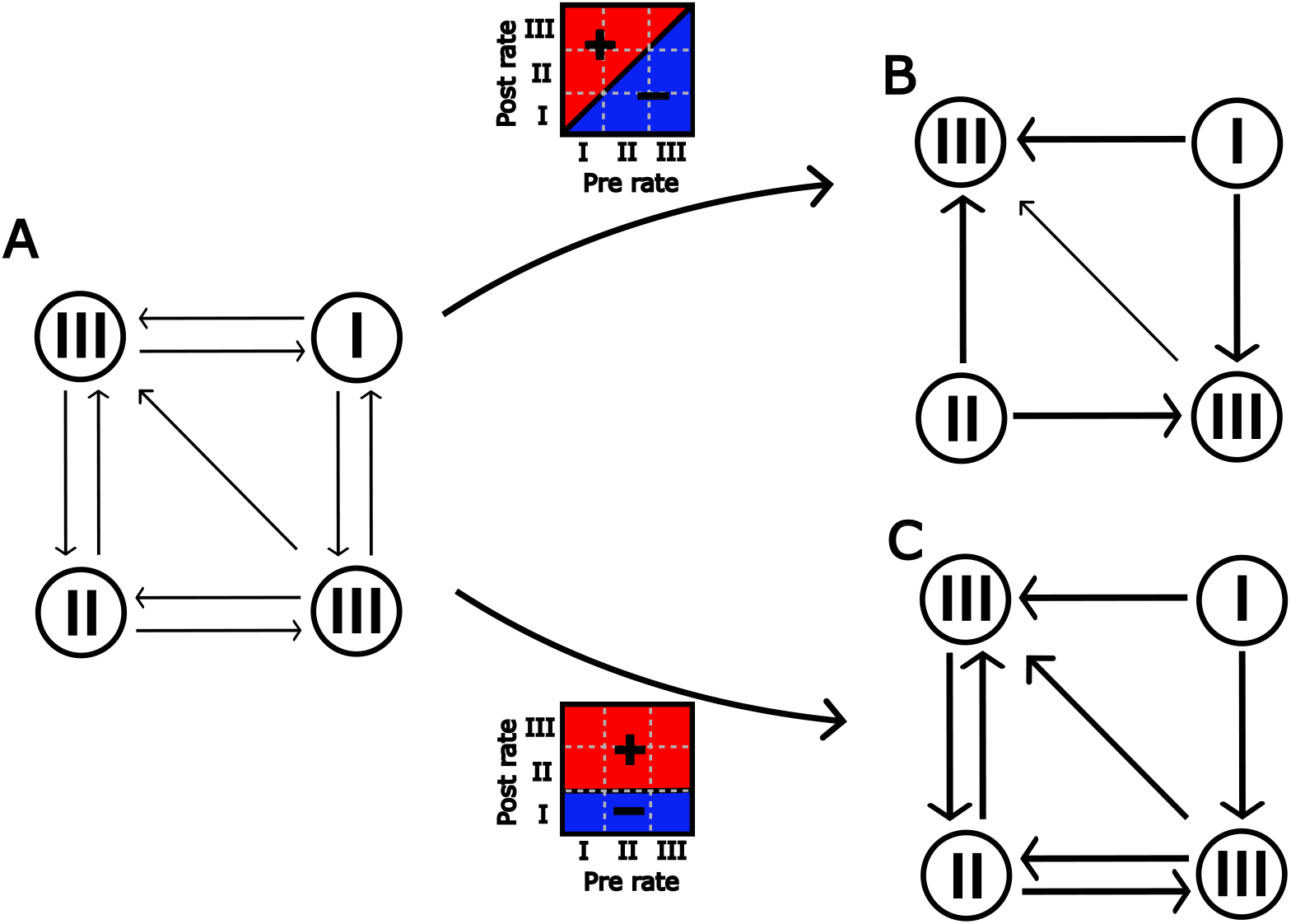
Simplified illustration of how the mean synaptic drift influences the emergence of sink and source nodes in RNNs. (**A**) A small RNN with four nodes of exhibiting firing rates from I to III. Updating the connection weights with a diagonally symmetric update rule with respect to the presynaptic and postsynaptic firing rates leads to the formation of sink and source nodes in (**B**). (**C**) While the non-symmetric update rule also morphs the most inactive nodes I into source nodes, the most active nodes III retain connections to II nodes.

Whether the Triplet and SL-STDP rules modify the connectivity in balanced RNNs [32] in a similar manner to that depicted in Fig 5 in practice is not obvious, since input-output correlations and the interplay between excitatory and inhibitory populations may produce unpredictable dynamics. To test how the SL-STDP model differs from the Triplet model, we perform network simulations and examine the changes in excitatory-to-excitatory (E-E) connections. We create spiking recurrent networks consisting of *N*_*e*_ = 800 excitatory and *N*_*i*_ = 200 inhibitory leaky integrate-and-fire neurons with a simple adaptive current. In the first network, E-E connections are modelled using the fitted SL-STDP synapse. In the second network, E-E connections are modelled using an uncoupled TM-Triplet model. The uncoupled TM-Triplet model consists of two components: the fitted Triplet model (see Methods) and the TM-model (Eq (1-3)). The TM-model parameters for the uncoupled TM-Triplet model are identical to those used in the SL-STDP model. In both networks, excitatory-to-inhibitory (E → I) connections are modelled using a depression-dominated TM synapse, i.e. the fitted SL-STDP synapse with *f* = 0, unless otherwise stated. See S6 for synapse models’ parameter comparison. Inhibitory-to-inhibitory and inhibitory-to-excitatory connections are static in both networks. We stimulate all neurons in both networks with Poisson spike trains. Before self-organization, the resulting networks exhibit balanced dynamics with a mean excitatory firing rate *λ*_exc_ = 39 Hz and a mean inhibitory firing rate *λ*_inh_ = 39 Hz. The coefficient of variation (CV) is approximately 0.3 for both populations suggesting some form of regularity in asynchronous firing times.

In order to study how the networks self-organize, we examine how the total E-E out-degree strength and the total E-E in-degree strength evolve as a function of the mean firing rate for each excitatory neuron. Fig 6 shows these distributions, along with the mean firing rate distributions from 10 seconds of self-organization to 30 minutes of self-organization. Interestingly, we find that the SL-STDP network exhibits a wide range of mean firing rates from 16 to 130 Hz and develops four peaks in the firing rate distribution (Fig 6A-B, right panel). This heterogeneity in mean firing rates is somewhat surprising, as the models in Fig 6 do not include any homeostatic control mechanism which is typically needed for balancing the synaptic growth of E-E synapses [3, 4, 33, 59]. As expected, in contrast to the SL-STDP network, Fig 6C-D shows how the uncoupled TM-Triplet network rapidly develops a unimodal firing rate distribution, with all the weights saturating near the maximum value *w*_max_, which was set to ten times the initial weight.

**Fig 6.**
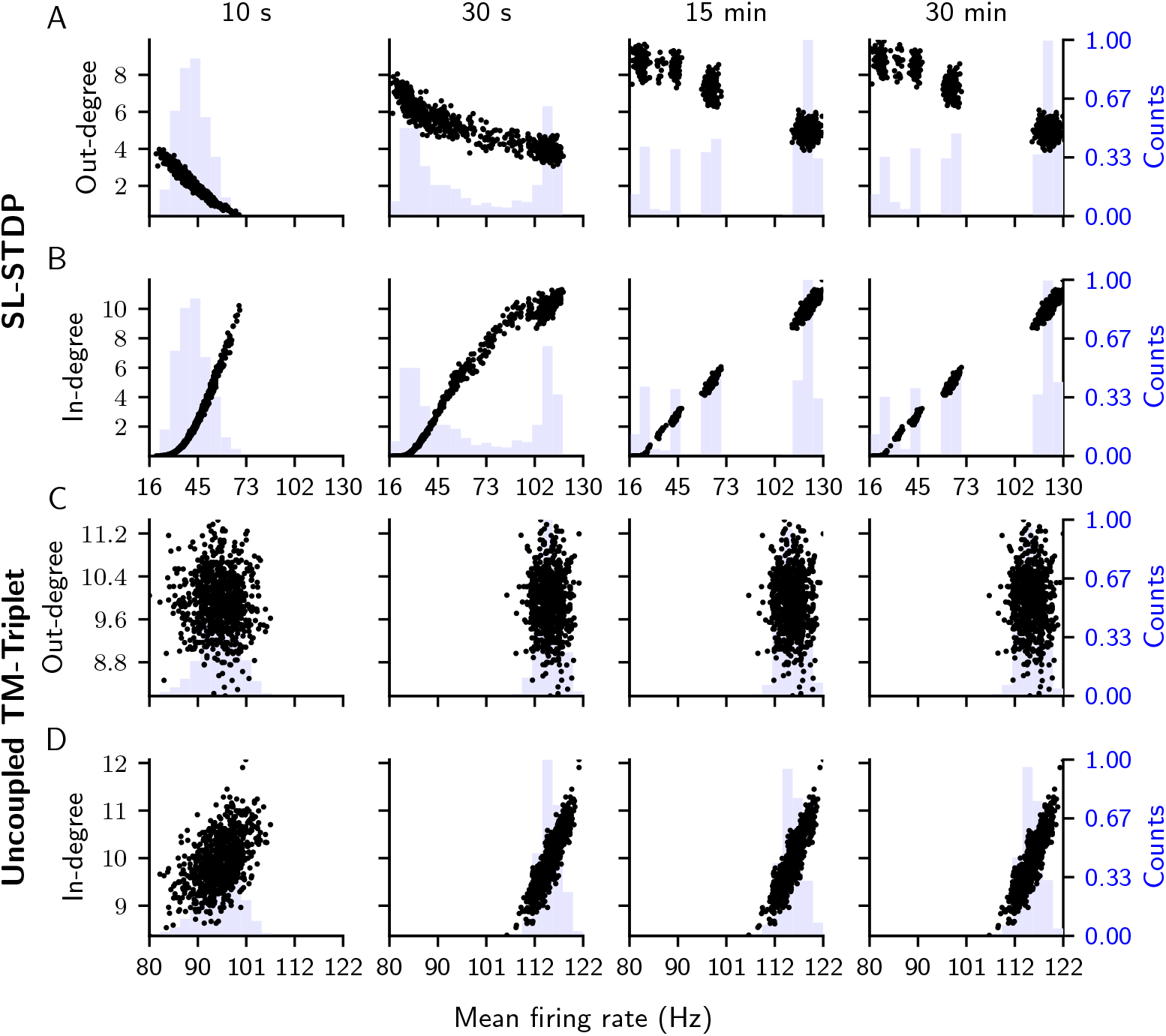
Evolution of relative total out-degree and in-degree strengths with respect to mean firing rates. Panels from left to right show the evolution of weight strengths from 10 seconds to 30 minutes of self-organization with respect to a mean firing rate of each excitatory neuron. Histograms of the mean firing rates are shown in light blue. (**A**) Total out-degree strength relative to the initial total out-degree for each of the *N*_*e*_ = 800 excitatory neurons in the SL-STDP network, shown as black dots. Each normalized total out-degree is plotted against the mean firing rate of the corresponding excitatory neuron. (**B**) Similar as (**A**), but for the relative total in-degree strength. (**C**-**D**) Relative total out- and in-degree strengths as functions of mean firing rate for the uncoupled TM-Triplet network.

Before self-organization, networks did not exhibit network-wide bursting. However, after self-organization, both networks display bursting activity, which can be attributed to the interplay between the adaptation current and the overall strengthening of the recurrent E-E weights [61]. To quantify the bursting activity, we estimate the bursting frequency and the total synchrony of the excitatory population by using the spike-contrast metric [62] after 30 minutes of self-organization. The maximal synchrony value *S* in spike-contrast metric is defined as the maximum synchrony over all bin sizes Δ. The synchrony itself is calculated as a product of the relative number of active neurons in a bin Δ, ActiveST, and the difference in the number of spikes in adjacent bins, i.e. Contrast [62].

As expected, the uncoupled TM-Triplet network exhibited excessive spiking during bursts, leading to exaggerated instantaneous firing rates (see Fig 2 in S1). The bursting frequency for the uncoupled TM-Triplet network was approximately 3 Hz, and the maximum synchrony was close to the theoretical maximum, *S*_Triplet_ = 0.93 with Δ = 145.4 ms. In contrast, the SL-STDP network displayed faster oscillations, with a frequency of 26 Hz and *S*_SL_ = 0.38 with Δ = 21.8 ms. The network activity of the SL-STDP network after 30 minutes of self-organization is shown in Fig 7A, and the spike-contrast metric for the excitatory population is shown in Fig 7B. We verified that the self-organization into four different mean firing rate peaks is indeed stable by simulating the SL-STDP network for 60 minutes. We observed that the four peaks persisted, and the bursting frequency remained at 26 Hz. Similarly, the maximum synchrony was *S*_SL_ = 0.39 with Δ = 19.6 ms, indicating the stability of the oscillations.

**Fig 7.**
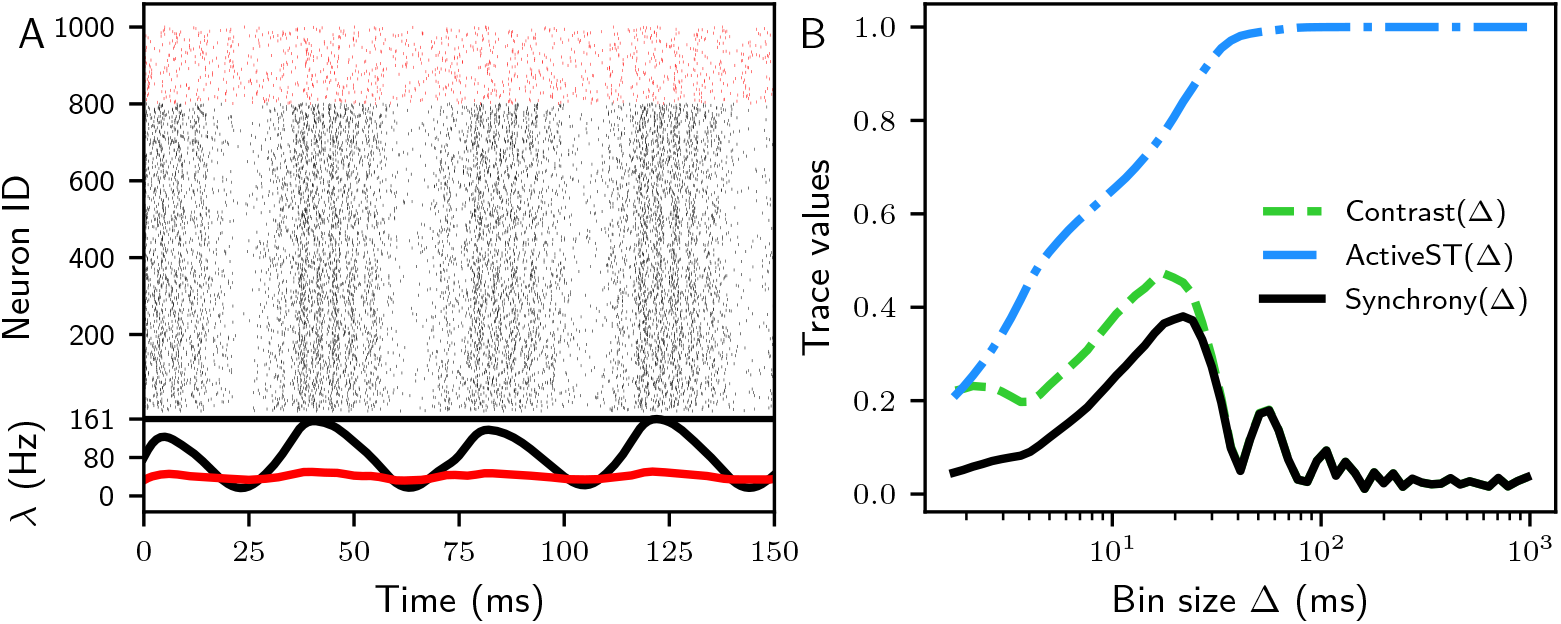
SL-STDP network’s activity after 30 minutes of self-organization exhibit network bursting and synchronization. (**A**) Raster plot of the SL-STDP network after 30 minutes of self-organization. Black markers show spike times of excitatory neurons, and red markers spike times of inhibitory neurons. At the bottom, the averaged population activity *λ* is shown for excitatory neurons (black) and inhibitory neurons (red). The excitatory population exhibits steady oscillations, whereas the inhibitory population remains asynchronized. (**B**) Spike-contrast metric [62] for the excitatory spikes shown in (**A**). Synchrony is measured as the product of Contrast and ActiveST (active spike trains). The maximum synchrony of 0.38 is achieved with a bin size Δ = 21.8 ms.

The mean firing rate of the excitatory population before self-organization *λ*_exc_ = 39 Hz is rarely observed *in vivo* for prolonged periods [63–65]. This motivated us to investigate how the SL-STDP model self-organizes under more realistic conditions. We found that the self-organization into distinct firing rate groups also occurs at lower levels of excitation but becomes more pronounced at higher levels of excitation. The same correlation trends between in- and out-degrees to mean firing rates are also preserved in lower mean firing rates. For more details, see S2.

### Homeostatic mechanisms shape connectivity in both systems

The expected runaway potentiation of E-E connections with the Triplet model is typically counteracted by homeostatic control mechanisms [3, 4, 33]. In the next set of simulations, we examine how the SL-STDP model and the uncoupled TM-Triplet model compare when such homeostatic mechanisms are applied. We begin with divisive weight normalization [33–36] and then investigate E → I plasticity that is less studied compared to inhibitory plasticity [37–41].

We apply weight normalization to excitatory in-degree weights every two seconds of simulation. Fig 8 shows the in-degree and out-degree strengths prior to applying weight normalization. Weight normalization limits the in-degree and out-degree strengths and the mean firing rates in both networks (Fig 8). The mean firing rates now exhibit unimodal distributions in both networks. The relationship between in-degree strength and firing rate is qualitatively similar across the two networks, although the uncoupled TM-Triplet network displays a larger overall scale. However, the relationships between out-degree strengths and mean firing rates differ substantially between the two models, as illustrated in Fig 8A and Fig 8C. As predicted in Fig 5B, the out-degree strength correlates negatively with the mean firing rate (see Fig 8A, Pearson correlation coefficient 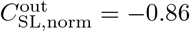 after 30 minutes of self-organization), while the in-degree strength correlates positively (Fig 8B, 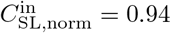) for the SL-STDP synapse. The same correlations are visible also for the non-normalized network (Fig 6A-B, 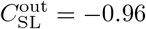 and 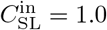. In contrast, the uncoupled TM-Triplet network with weight normalization exhibits a positive correlation between out-degree strength and mean firing rate (Fig 8C, 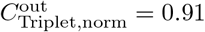).

**Fig 8.**
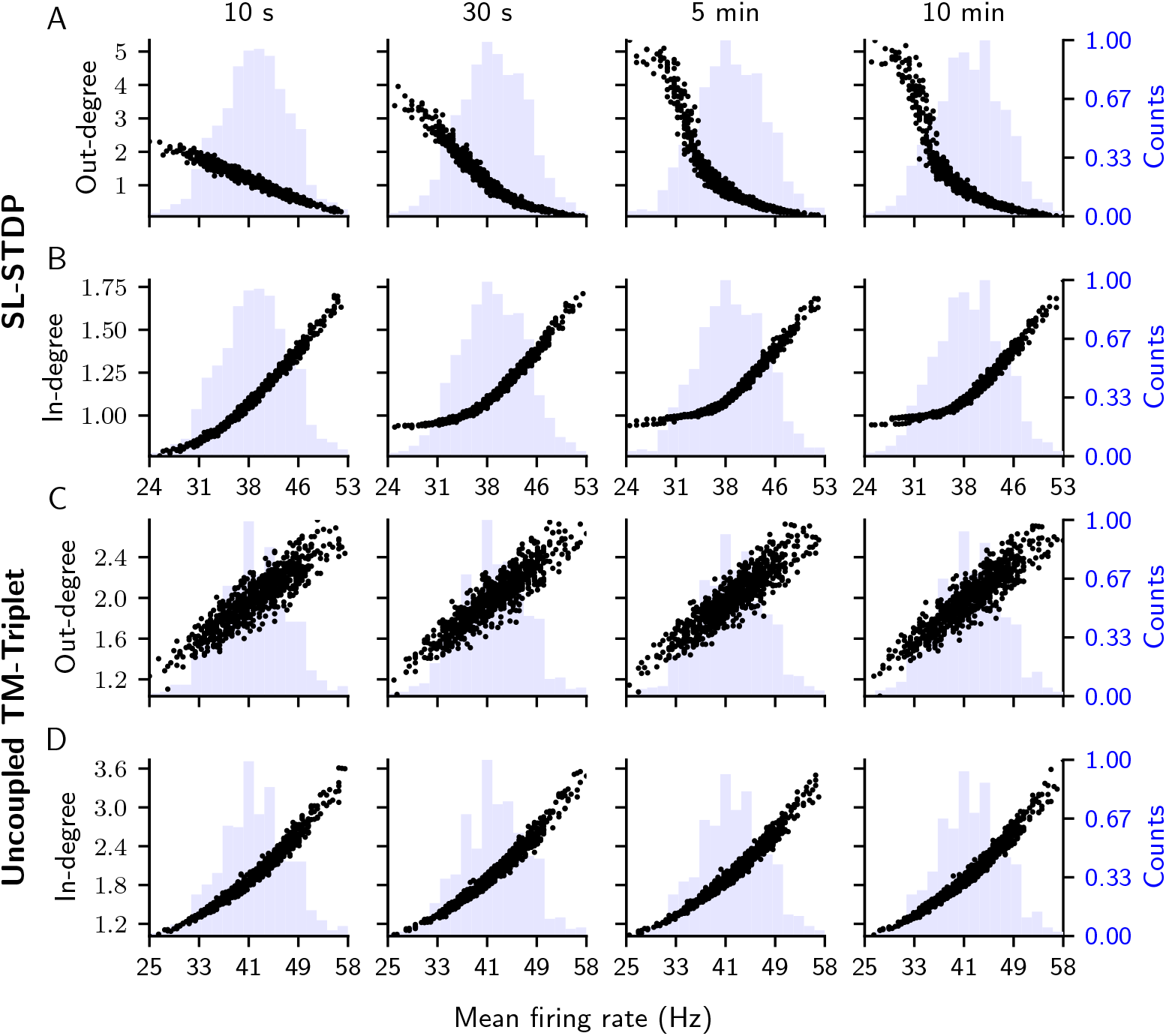
Evolution of the relative total out-degree and in-degree strengths with respect to mean firing rates in weight-normalized networks. Panels from left to right show the evolution of weight strengths from 10 seconds to 10 minutes of self-organization with respect to the mean firing rate of each excitatory neuron. Histograms of the mean firing rates are shown in light blue. In-degree weights are normalized every two seconds. The in-degree weights are shown immediately before the normalization step. (**A**) Total out-degree strength relative to the initial total out-degree for each of the *N*_*e*_ = 800 excitatory neurons in the SL-STDP network, shown as black dots. Each normalized total out-degree is plotted against the mean firing rate of the same neuron. (**B**) Similar as (**A**), but for the relative total in-degree strength. (**C**-**D**) Relative total out- and in-degree strengths as functions of the mean firing rate for the uncoupled TM-Triplet network.

Most notably, networks with the weight normalization do not produce bursting activity in contrast to the non-normalized case (*S*_SL,norm_ = 0.06 and *S*_Triplet,norm_ = 0.05). Instead, the networks maintain asynchronous activity with a mean CV of 0.3. This can be attributed to the smaller scale of potentiated E-E weights resulting from weight normalization.

Next, we investigate how E → I plasticity shapes connectivity and network activity in the SL-STDP network by setting *f* = 1 for the E → I connections, meaning that the exact same plasticity model is applied to both E-E and E → I connections. In contrast to weight normalization, we find that the E → I plasticity preserves the emergence of mean firing rate clusters (Fig 9A-B). The E → I plasticity promotes faster network oscillations (50 Hz compared to 26 Hz), which is also reflected in a shift in the excitatory population’s maximal synchronization (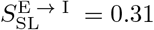 with Δ = 10.4 ms compared to *S*_SL_ = 0.38 with Δ = 21.8 ms). Interestingly, the previously asynchronous inhibitory population (Fig 7A), also starts to oscillate as a result of E → I plasticity, with the same frequency of 50 Hz. This suggests that the E → I connection strength is net increasing, reaching a level sufficient to drive the inhibitory population into an oscillatory regime. See S3 for more detailed inhibitory activity analysis.

**Fig 9.**
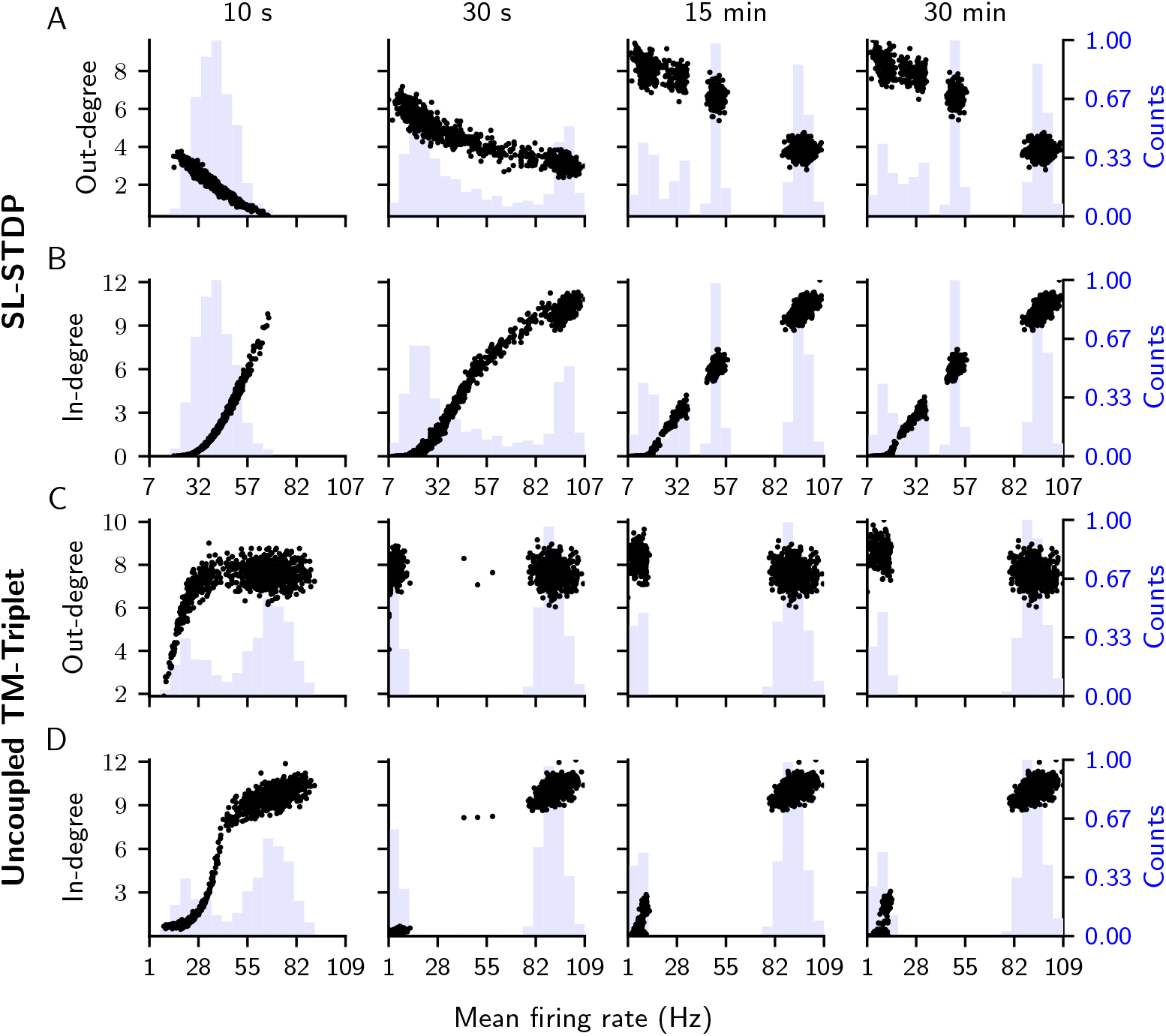
Evolution of the relative total out-degree and in-degree strengths with respect to mean firing rates in networks with E → I plasticity. Both models include E-E and E → I plasticity and no weight normalization. Panels from left to right show the evolution of weight strengths from 10 seconds to 30 minutes of self-organization with respect to the mean firing rate of each excitatory neuron. Histograms of the mean firing rates are shown in light blue. (**A**) Total out-degree strength relative to the initial total out-degree for each of the *N*_*e*_ = 800 excitatory neurons in the SL-STDP network, shown as black dots. Each normalized total out-degree is plotted against the mean firing rate of the corresponding neuron. (**B**) Similar as (**A**), but for the relative total in-degree strength. (**C**-**D**) Relative total out- and in-degree strengths as functions of mean firing rate for the uncoupled TM-Triplet network.

Fig 10 elaborates the differences in the clustering of SL-STDP networks with (Fig 10C-D) and without (Fig 10A-B) E → I plasticity after 30 minutes of self-organization. In both cases, we observe five different clusters C1 to C5 with varying sizes (Fig 10A and C). E → I plasticity decreases the size of the cluster C5 (*n* = 310 with vs. *n* = 400 without) while increasing the number of neurons in other clusters, except C3. Number of connections between the clusters are shown in Fig 10B for the network without E → I plasticity and in Fig 10D for the network with E → I plasticity. Both matrices are close to triangular indicating a strong directionality in the network organization. Indeed, both SL-STDP networks self-organize to have mostly feed-forward connections, as shown in Fig 10E for the SL-STDP network without E → I plasticity.

**Fig 10.**
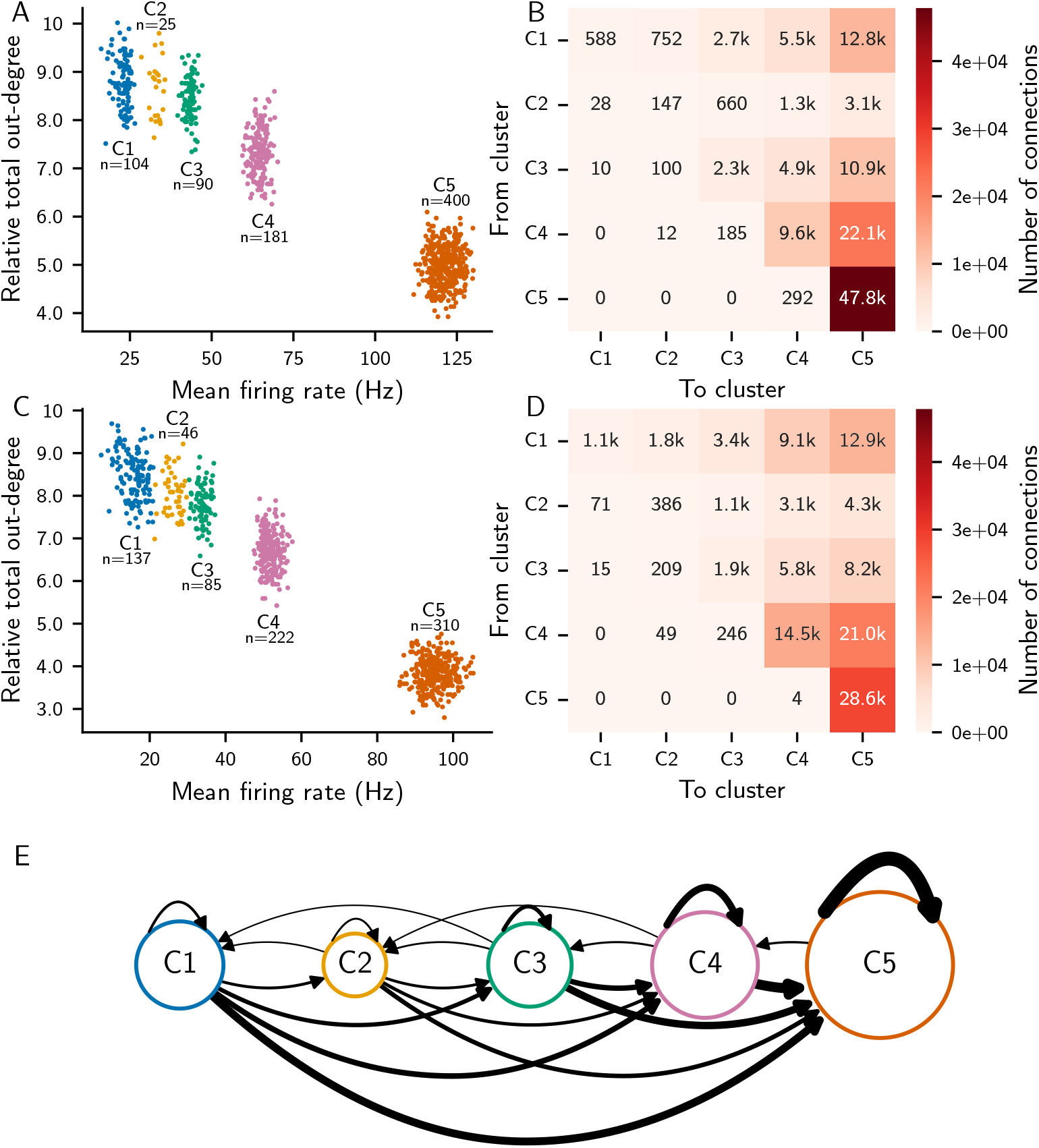
SL-STDP networks self-organize into separate clusters. (**A**) The SL-STDP network’s relative total out-degree with respect to the mean firing rate for each excitatory neuron after 30 minutes of self-organization. Essentially, the same data is shown as the rightmost panel in Fig 6A. Five clusters C1 to C5 are clearly visible. (**B**) Number of connections from each cluster to other clusters for the network in (**A**). Connections with small postsynaptic weights are not counted. (**C-D**) Same as in (**A-B**) but for the SL-STDP network with E → I plasticity, i.e. Fig 9A rightmost panel. (**E**) Visualization of how the network’s clusters in (**A**) are connected. Size of the nodes C1 to C5 refer to a population size *n* and the width of the arrows reflect the number of connections between the clusters.

The Triplet network undergoes more drastic changes in in- and out-degree distributions when E → I plasticity is included. Instead of forming a single hyperactive group of neurons, two clusters emerge (Fig 9C-D). Surprisingly, there is little separation in out-degree strengths between the two clusters, although the in-degree strengths are highly polarized (Fig 9C-D). Taken together, these results demonstrate that direct coupling between short-term dynamics and long-term plasticity produces connectivity and network activity that differ fundamentally from traditional approaches, in which short-term dynamics do not directly influence long-term plasticity.

### SL-STDP model improves information capacity in RNNs

We extend the comparison between the systems to their information capacities to assess whether the changes in connectivity have practical implications for the information processing capabilities of the networks. To quantify the working memory capabilities of the networks, we evaluate their performance on tasks of varying difficulty. Specifically, we employ firing rate encoded fading memory [43] tasks with increasingly challenging nonlinear input-output transformations [42, 45].

The workflow for this task is as follows: 1) The network is allowed to self-organize for 30 minutes (except 10 minutes with the weight normalization condition) with only background noise. During this step, long-term plasticity is active. 2) Long-term plasticity is then turned off, and the network is stimulated with an input signal (Poisson process with a stepwise changing rate). 3) The system is simulated for X steps, and a linear readout is fitted to the neural activity data. 4) Stimulation is continued with a new input for Y steps, and the fitted linear readout is used to predict the output. The capacity of the network is calculated by comparing the predicted output with the ground truth, which is defined as a delayed nonlinear transformation of the input signal.

There are two varying components that determine the difficulty of the task assigned to the network. First, the output target is delayed in time from zero to three input steps, zero meaning that the network should perform the input-output transformation immediately and non-zero values meaning that the network has to memorize the input for a given time. Secondly, the output target is a nonlinear transformation of the input signal defined by Legendre polynomials. Increasing the total polynomial degree results in a more difficult signal transformation [42]. See Methods for details. Simultaneously varying of these two components yields a 4-by-4 matrix, which we use for the capacity evaluation. The capacities reported in Fig 11 and Fig 12 are normalized by the capacity values of a network without any long-term plasticity rule. As such, positive values in Fig 11 and Fig 12 indicate that a network has better information capacity than the network without self-organization and negative values indicate that a network has worse information capacity compared to the network without self-organization. The absolute capacity values of the baseline network are shown in S4.

**Fig 11.**
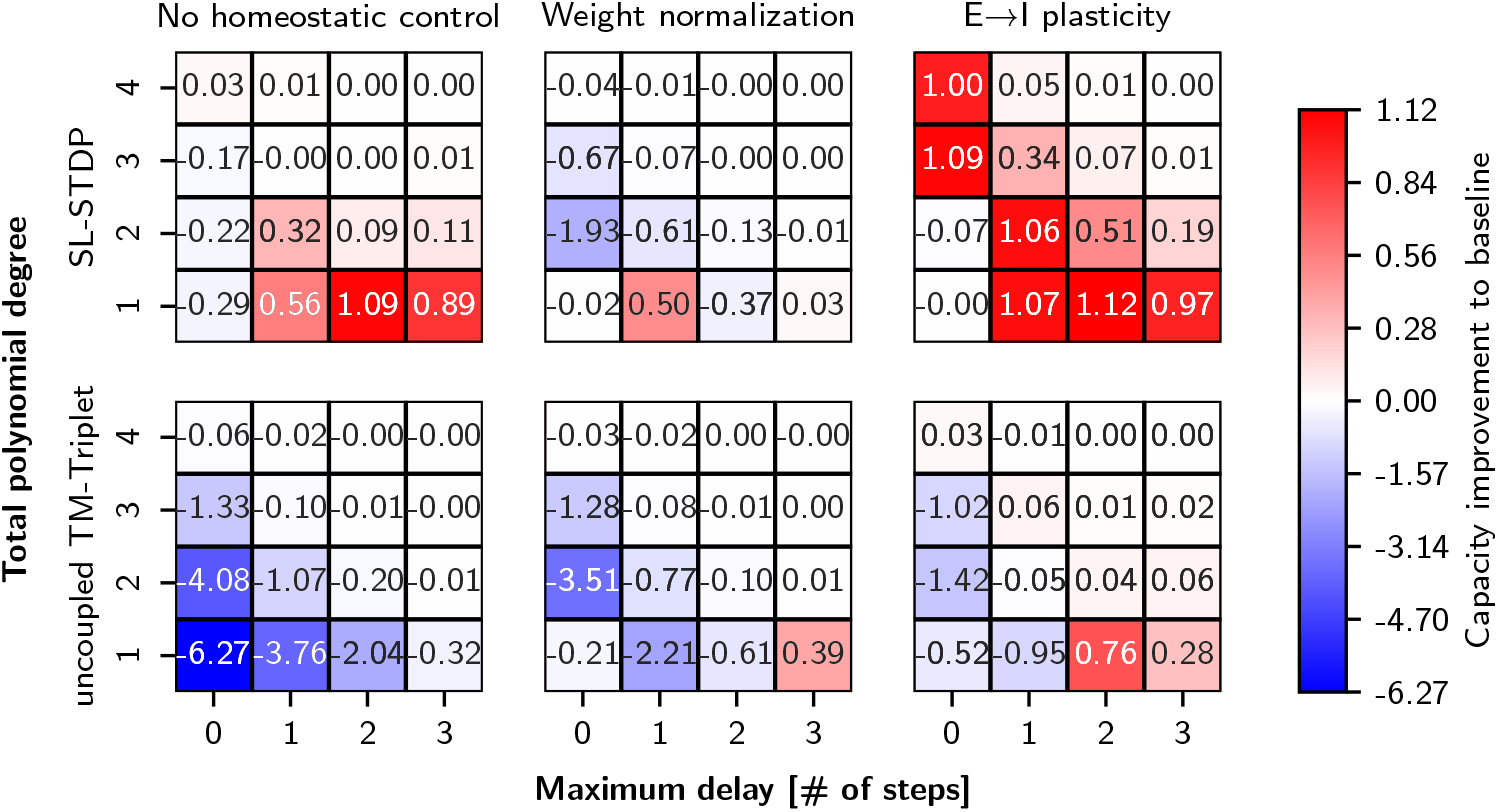
Information capacities for the SL-STDP and the uncoupled TM-Triplet models under different homeostatic conditions. Reported capacities are normalized by the capacity values of an RNN that has only short-term dynamics. For the absolute capacity values of the baseline network, see S4. Higher values indicate better performance in output prediction. Increasing the total polynomial degree and maximum delay corresponds to more difficult working memory tasks. The reported numbers are mean testing capacities averaged over ten different random initializations of the networks. See S4 for the standard deviations.

**Fig 12.**
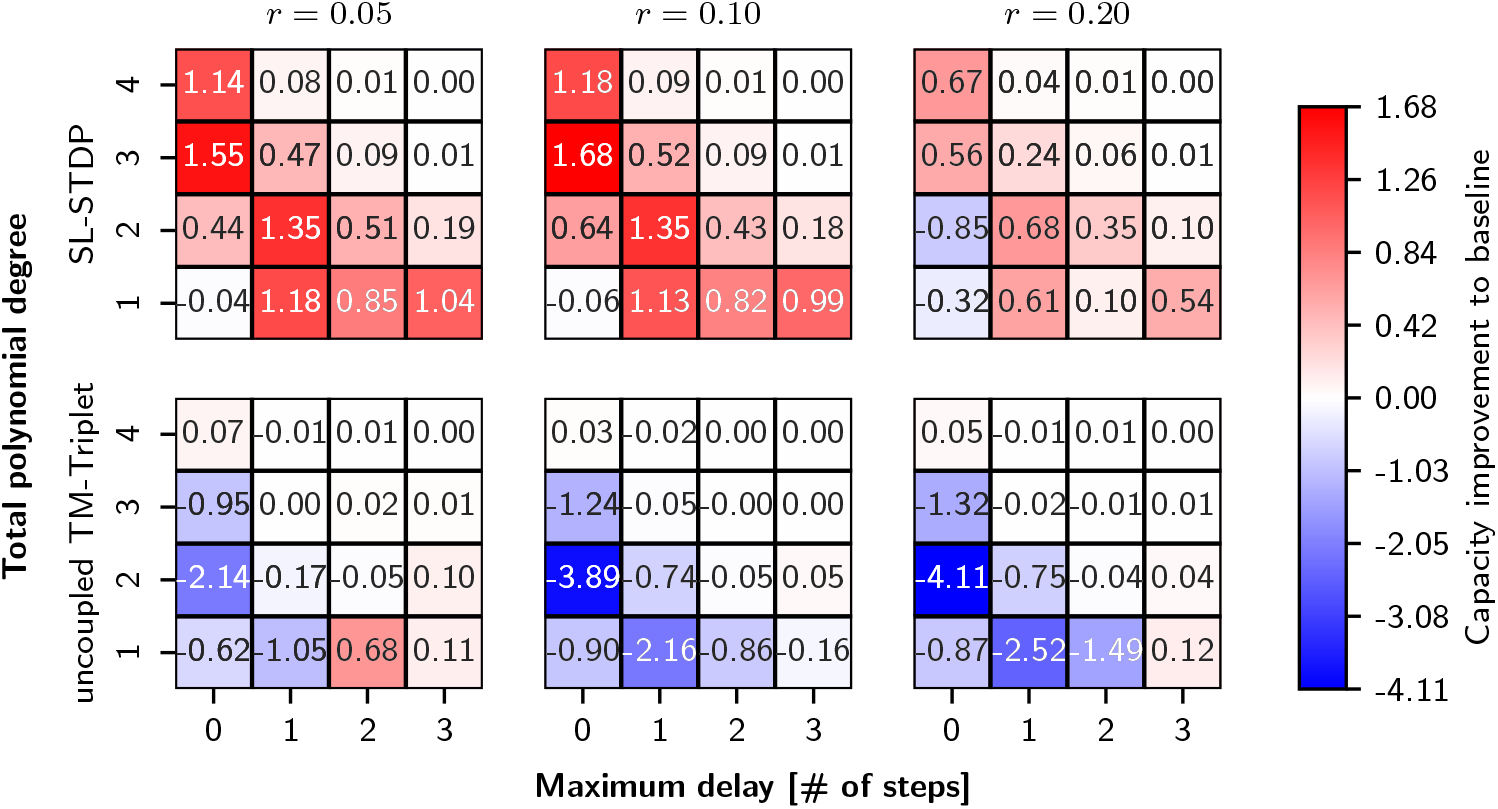
Information capacities for the SL-STDP and the uncoupled TM-Triplet models with varying ratios of facilitating synapses. The ratio *r* denotes the ratio between facilitating and depressing E → I synapses. All E → I connections are plastic. Reported capacities are normalized by the capacity values of an RNN that has only short-term dynamics. For the absolute capacity values of the baseline network, see S4. Higher values indicate better performance in output prediction. Increasing the total polynomial degree and maximum delay corresponds to more difficult working memory tasks. The reported numbers are mean testing capacities averaged over ten different random initializations of the networks. See S4 for the standard deviations.

Fig 11 shows information capacities for all the networks studied in the previous sections. All models, including the network without long-term plasticity (see S4), perform poorly on tasks requiring both complex nonlinear transformation and long memory (the 2×2 submatrix in the top-right corner). However, for all other combinations, we observe varying performance. As expected, the uncoupled TM-Triplet model without homeostatic control performs poorly on all tasks. Both weight normalization and E → I plasticity restore the network’s basic functionality to some extend, while E → I plasticity outperforms weight normalization in most of the tasks.

Interestingly, the SL-STDP model shows better linear memory compared to the baseline network, even without homeostatic control. We associate this to the high variety of in- and out-degree values (Fig 6B, last panel). Overall, weight normalization reduces the capacity of the SL-STDP model. One possible explanation for this could be the absence of neurons with high in-degree strength (Fig 8A-B, last panel) that could integrate signals from other nodes. In contrast, the E → I plasticity improves capacities in almost all tasks in the SL-STDP network. The simultaneous improvement of linear memory and nonlinear computation is encouraging, since a trade-off between memory and nonlinearity is reported for many reservoir computing systems [42, 45, 66].

The performance of E → I plasticity prompted a more detailed investigation of the effects of E → I plasticity on network capacity. In Fig 11, all the E → I connections are modeled similarly as the E-E connections. However, there is a body of evidence that also facilitating synapses exist between pyramidal cells and interneurons [47, 67–69].

Therefore, we perform experiments using E → I plasticity with a varying ratio *r* between facilitating and depressing synapses. The facilitating connections are modeled using the parameters shown in Table 1 (row ‘Facilitation’). All E-E connections remain unchanged. The results are shown in Fig 12 for both the SL-STDP network and for the uncoupled TM-Triplet network.

We found that increasing the number of facilitating connections is beneficial for the SL-STDP model while the performance of the uncoupled TM-Triplet model decreases.

The SL-STDP network achieves better performance at lower ratios *r* = 0.05 and *r* = 0.10 compared to *r* = 0.20 across all capacity tasks (Fig 12). Since firing rate encoded information capacity tests are sensitive to the overall network excitability, we measured the mean inhibitory and excitatory activities during the capacity computation. We observed increasing inhibitory activity trend with more facilitating synapses in the SL-STDP network model. This is not surprising, as facilitating SL-STDP synapses potentiate more rapidly (Fig 1), thereby driving stronger inhibitory activity. In response, the average excitatory rate first decreased from 33 Hz to 20 Hz but then started to increase after *r* > 0.1. In contrast, the uncoupled TM-Triplet model shows high inhibitory activity across all values of *r*, but is still unable to stabilize excitatory firing rates, resulting in a reduced total information capacity, as seen in Fig 12.

We also performed information capacity tests with an uncoupled TM-Triplet model in which long-term plasticity parameters were not fully constrained by a biological data. In this case, the network had slower runaway activity and E → I plasticity was able to stabilize the excitatory rates. Information capacities for this biologically unconstrained uncoupled TM-Triplet model were significantly higher than that of the data fitted uncoupled TM-Triplet model, but still underperformed compared to the SL-STDP model. All together, we find that an RNN that employs the SL-STDP synapse model for E-E and E → I connections (with a small number of facilitating synapses, *r* = 0.1) during self-organization achieves the best information capacities on the reservoir computing tasks.

The two key differences between the SL-STDP model and the uncoupled TM-Triplet model are the sharing of the latent state between short-term dynamics and long-term plasticity and the inclusion of gain control on the postsynaptic side in the SL-STDP model. This raises an important question: is sharing the state between short-term dynamics and long-term plasticity alone sufficient to reproduce superior information capacities? To address this, we repeated the model fitting, analysis, and the working memory test for a variant of the SL-STDP model that does not include postsynaptic gain control. We found out that the main results hold also for this model variant, indicating that the balance between the LTP and LTD regimes is the most important factor underlying the improved information capacity. See S5 for details.

## Discussion

Integrated models of short-term and long-term plasticity have recently gained increasing attention [19, 22, 24], but their effects on network dynamics and information capacity in RNNs have not been studied systemically. For this purpose, we developed a phenomenological and intuitive model of unified short-term dynamics and postsynaptic long-term plasticity by taking components from the Tsodyks-Markram (TM) model [27, 30] and the Triplet model [8]. We fitted the model to pyramidal neuron data from the rat layer 5 visual cortex [31] and investigated the effects on network connectivity and information capacity in RNNs, both with and without homeostatic control mechanisms [33]. For comparison, we repeated the same fitting procedure and network analyses on the uncoupled TM-Triplet model. We found that the network with the SL-STDP synapse model exhibits richer information capacity compared to the network with the uncoupled short-term dynamics and long-term plasticity.

We studied the mean synaptic weight drift of the SL-STDP model in order to understand the impact of coupling short-term and long-term plasticity. Interestingly, we observed a stark contrast in the mean synaptic weight drifts between the SL-STDP and Triplet models, although both models were fitted to the same data set [31] (Fig 4). The difference in the drifts stems from the sparsity of experimental data points, which lie only along the diagonal of the frequency dependency matrix. To our knowledge, no electrophysiological data set systematically quantifying the impact of independently varying pre- and postsynaptic firing rates is currently available, thus posing a challenge for the model validation. In studies of rodent cortex [70–72], it is found that neurons with facilitating synapses have a preference for reciprocal connections, whereas neurons with depressing synapses predominantly exhibit unidirectional connections. Such connection motifs appear naturally with the SL-STDP synapse model, since short-term depression promotes symmetric frequency dependency, while short-term facilitation promotes a Triplet-like fixed threshold for the potentiation, as seen in Fig 1. In conclusion, our model predicts that the frequency dependence of long-term plasticity is modified by short-term plasticity, thereby explaining the correlation between connectivity motifs and short-term dynamics [70–72]. However, the precise form remains to be validated by future studies.

To examine how short-term dynamics induced frequency dependence of long-term plasticity affects network activity and connectivity, we applied the SL-STDP synapse model to an RNN consisting of adaptive leaky integrate-and-fire excitatory and inhibitory neurons. The network was stimulated with homogeneous Poisson processes, and we observed how the network self-organizes under this input. As anticipated from the mean synaptic weight drift (see Fig 4B and Fig 5), we observed a positive correlation between in-degree strength and firing rate and a negative correlation between out-degree strength and firing rate in Fig 6A-B, implying an anti-correlation between in-degree and out-degree strengths. Anti-correlations between in-degree and out-degree strengths in RNNs have been studied before and have been found to be beneficial for stimulus detection and protecting against pathological bursting [73], which aligns well with our results. Per-neuron in-degree and out-degree correlations are commonly less studied in experimental settings, but recent large-scale connectomics studies may help to address this gap [74, 75].

The interplay between the TM model and the Triplet STDP model in RNNs has previously been examined without direct coupling between the mechanisms [76]. Similar findings regarding the emergence of connectivity motifs involving the interaction of short-term dynamics and long-term plasticity have been reported, although the underlying mechanism differs from that described here. In the study by Vasilaki and Giugliano [76], facilitating synapses increased neuronal firing rates into the net LTP regime, thereby promoting the formation of reciprocal connections. In contrast, neurons with depressing synapses exhibited firing rates below a critical threshold, where spike timing plays a more significant role than rate dependency. In this region, unidirectional connections dominate [76]. As a consequence, neurons with depressing synapses can also exhibit reciprocal connectivity motifs if the external drive is strong enough. We demonstrated this indirectly in Fig 6C-D, where a highly interconnected homogeneous population emerges even with depressing short-term synapses. In contrast, the network with SL-STDP synapses forms clusters of neurons that operate at different firing rate ranges (Fig 6A-B and Fig 10). Thus, the modification of the frequency dependence of long-term plasticity by short-term dynamics is fundamentally different from the uncoupled version [76], in which short-term dynamics influence only spiking activity and do not directly affect the long-term plasticity of the neurons.

The clusters emerged in various scenarios, highlighting the stability of this emergent property. The self-organized clusters exhibit a clear feed-forward structure as shown in Fig 10E. As a comparison to synfire chains [77], clusters of neurons have skip connections in addition to sequential connections (Fig 10E). We found that the input scheme with a lower level of excitation affects clustering by reducing the number of neurons in higher firing rate clusters, but still three different well-separated clusters appeared (see S2). On the other hand, removing postsynaptic gain control revealed that the increased maximal firing rate generated one more cluster (see S5). Thus, there seems to be a correlation between the size and number of clusters and network excitability. Removing the adaptive current from the neuron model eliminated network oscillations but did not completely abolish the emergence of clusters, indicating that the emerging network-wide oscillatory dynamics are not required for clustering to occur.

Weight normalization [34–36] prevented the emergence of the clusters (Fig 8A-B), consistent with previous studies showing that afferent synaptic normalization leads to unimodal firing rate distributions (see e.g. [78]). Nevertheless, weight normalization preserved the anti-correlation between out-degrees and mean firing rates for the SL-STDP model as a key difference to the uncoupled TM-Triplet model. The divisive weight normalization applied at fixed time intervals is rather abrupt and global constraint that may be non-biological, as discussed in [79]. It would be interesting to investigate how the SL-STDP model differs from the uncoupled TM-Triplet model in existence of more detailed weight normalization rules. Further analysis with homeostatic balancing mechanisms revealed that the clusters were preserved with E → I plasticity [37–41] (Fig 9A-B). E → I plasticity affected the size of the clusters and the connectivity between the clusters, as shown in Fig 10. Another form of homeostatic control mechanism that is commonly applied to RNNs is inhibitory plasticity (see e.g. [3, 80, 81]). Investigating how the SL-STDP synapse model interacts with inhibitory plasticity would be an interesting topic for future research.

We did not anticipate the emerge of clusters, and we currently lack a descriptive theory to explain this phenomenon. Further studies incorporating, for example, LTD update weight dependency [82, 83] or a BCM-like adaptive threshold [55], could help to clarify the mechanisms that contribute to the emergence of clusters. During the early phases of self-organization (Fig 6A-B), firing rates exhibit a skewed unimodal distribution resembling a lognormal distribution. Lognormal firing rate distributions are commonly observed in many brain regions (for review, see [84]). It is rather interesting whether the clustering might occur during the brain development or in conditions where excitation-inhibition balance is impaired.

Finally, we quantified how the learned connectivity affects the working memory capabilities of the networks by performing a set of information capacity tests with varying levels of difficulty [42]. We found that the networks incorporating the SL-STDP synapse model perform well even without any additional homeostatic control mechanisms, while the uncoupled TM-Triplet model performs poorly as expected (Fig 11). Including weight normalization did not improve performance in the SL-STDP network, indicating that the clustering during high-input self-organization is useful for the network’s information processing capabilities. In contrast to weight normalization, E → I plasticity showed improved capacity for both fading memory and nonlinear computation. Furthermore, changing a fraction of the E → I connections from depressing to facilitating synapses, suggested by the heterogeneity between excitatory and inhibitory connections [67, 68], improved information capacity even further (Fig 12). The role of heterogeneity in short-term plasticity parameters, from the perspective of information processing, is also supported by a recent study [85], in which the Tempotron rule [86] is used to modify short-term plasticity parameters. Our model suggests that such heterogeneity in presynaptic short-term dynamics also influences postsynaptic long-term changes. Further studies are needed for understanding how simultaneous presynaptic and postsynaptic long-term changes interact between each other.

The enhanced information capacity highlights that the networks trained with the SL-STDP rule show better noise robustness and can retain information longer for more demanding nonlinear transformations. The higher information capacity of the networks is particularly relevant for neuromorphic computing applications [87]. We hope that our minimal yet biologically grounded approach, which directly couples short-term and long-term plasticity, will prove useful for such applications in the future.

## Methods

### Minimal all-to-all Triplet model

The minimal all-to-all Triplet model is defined with one presynaptic state *c*(*t*) and two postsynaptic states *z*_1_(*t*) and *z*_2_(*t*) [8]:

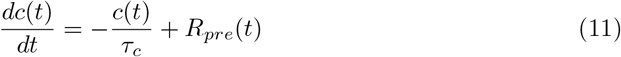

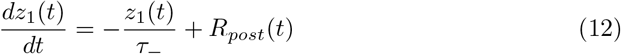

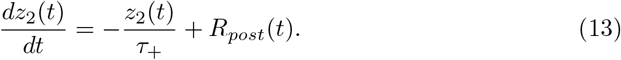

The time constant *τ*_*c*_ defines the timescale of the presynaptic trace, while *τ*_−_ and *τ*_+_ define the timescales of the postsynaptic traces. As for the SL-STDP model, *R*_*pre*_(*t*) is the presynaptic neural response function and *R*_*post*_(*t*) is the postsynaptic neural response function.

The LTD and LTP updates are then defined as

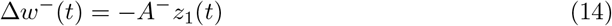

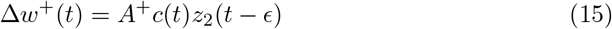

similarly to the SL-STDP model Eqs (6-7). The weight *w* is updated in the same manner as in Eqs (8-9).

The mean drift of *w* for the Triplet model is derived by assuming uncorrelated presynaptic and postsynaptic Poisson processes [8]:

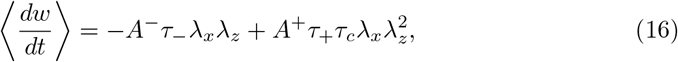

where *λ*_*x*_ is the firing rate for presynaptic spikes and *λ*_*z*_ is the firing rate for postsynaptic spikes. For a more detailed analysis with correlated inputs, see [88]. Eq (16) is a net positive when *λ*_*z*_ exceeds a threshold

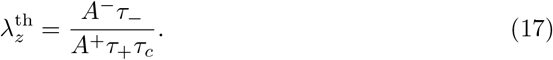

By default, 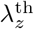 is constant but can be made adaptive in order to align with the BCM theory [8, 55].

### Numerical methods

#### Model fitting

The SL-STDP synapse model and the minimal all-to-all Triplet model are solved in an event-based manner (see e.g. [24, 47]) and fitted to pyramidal neuron data from the rat L5 visual cortex [31]. Three long-term plasticity induction data sets from [31] were included: 1) Δ*t* = *±* 10 ms pre-post spike interval with varying pre-pre and post-post spike time intervals, 2) stochastic data set, where the spike times were generated from a Gaussian distribution with a mean *µ*(Δ*t*) = 0 ms and a standard deviation *σ*(Δ*t*) = 7 ms and 3) a few additional data points with varying Δ*t* (Fig 7C in [31]). The fitting error *E*_*i*_ for each of the cases from 1) to 3) is calculated as the mean-squared error between the model’s value *w*^mod^ and the experimental value *w*^exp^:

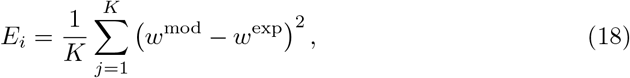

where *K* is the number of data points in each data set *i. w*^mod^ and *w*^exp^ are given as a percentage change from the initial synapse strength. We generated 20 different realization for each data point in the stochastic data set to reduce trial-to-trial variability. The total error is then defined as

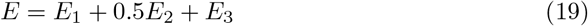

which is a subject of the minimization.

Error minimization is performed using the bounded limited memory

Broyden-Fletcher-Goldfarb-Shanno (L-BFGS-B) algorithm [89, 90]. In some cases, we observed that a few repetitive runs of the L-BFGS-B algorithm with basinhopping [91] resulted in a lower value of *E* than a single run of the L-BFGS-B algorithm. However, increasing the number of basinhops did not significantly reduce *E*, and thus the number of basinhops was set to five. The boundary conditions for the acceptable parameter regimes were selected by following reasonable biological values. For the exact boundary conditions, see https://github.com/IiroAhokainen/SL-STDP. The fitted parameter values are shown in Table 2 for the SL-STDP model and in S1 for the Triplet model. Also, see S6 for a parameter comparison.

#### Semi-analytical solution of nullclines

Nullclines for Fig 1 are obtained by calculating the partial derivatives of Eq (10) and setting them to zero:

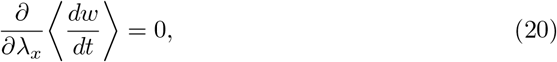

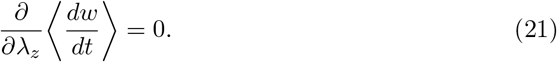

Both Eq (20) and Eq (21) yield to quartic functions of the form

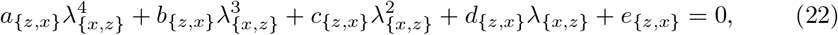

where multipliers *a*_*{z,x}*_, *b*_*{z,x}*_, *c*_*{z,x}*_ *d*_*{z,x}*_ and *e*_*{z,x}*_ are dependent on the model parameters and either *λ*_*z*_ or *λ*_*x*_. For the definitions of model parameters, see the Results section. For Eq (20), we find

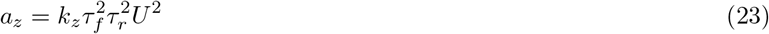

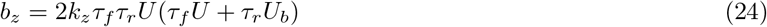

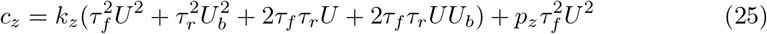

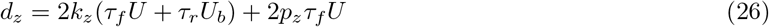

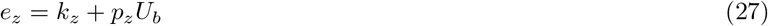

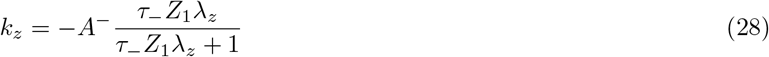

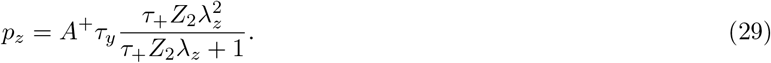

Similarly, for Eq (21), we find

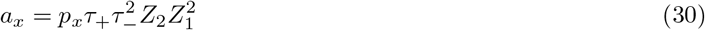

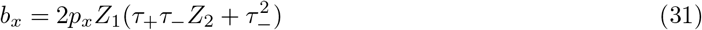

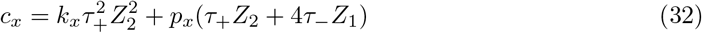

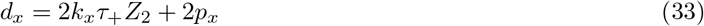

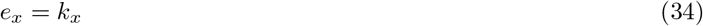

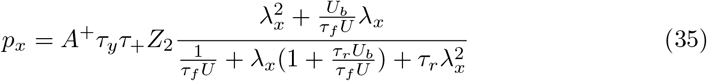

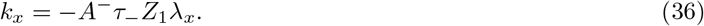

The quartic functions in Eq (22) were then solved numerically using the SymPy Python library [92], and only real positive roots were accepted as solutions. We ensured that the nullcline in Eq (20) was indeed giving local maxima by calculating the second partial derivative with respect to *λ*_*x*_. This yielded negative terms from 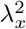 up to a polynomial 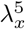, while only one polynomial *λ*_*x*_ and a few constants were positive.

### Neuron model

The neuron model used for all network simulations is a leaky integrate-and-fire neuron with a simple adaptation current *I*^adap^(*t*):

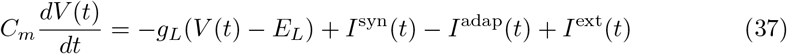

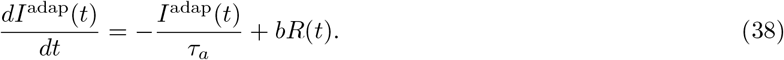

Here, *V* (*t*) is the membrane voltage, *C*_*m*_ is the membrane capacitance, *g*_*L*_ is the leak conductance, *E*_*L*_ is the leak reversal potential, *I*^syn^(*t*) is the synaptic current from other neurons, *I*^ext^(*t*) is the external current, *τ*_*a*_ is the time constant for the adaptation current. Lastly *b* is the action potential triggered step increase that drives the adaptation current to positive values. When *V* (*t*) reaches a spiking threshold *θ*, a spike is emitted and the neuron enters a refractory period for a time *t*_*ref*_, during which *V* (*t*) is clamped to a reset potential *V*_*reset*_ before continuing integration. All parameter values are identical for excitatory and inhibitory neurons and are listed in Table 3.

**Table 3.** Parameter values for the leaky integrate-and-fire neuron with adaptation current.

| Symbol | Definition | Units | Value |
| --- | --- | --- | --- |
| $C_m$ | Membrane capacitance | pF | 250 |
| $g_L$ | Leak conductance | nS | 25 |
| $E_L$ | Leak reversal potential | mV | -70 |
| $\tau_a$ | Adaptation time constant | ms | 150 |
| $b$ | Adaptation increment per spike | pA | 50 |
| $\theta$ | Spiking threshold | mV | -55 |
| $V_{reset}$ | Membrane reset potential | mV | -70 |
| $t_{ref}$ | Refractory period | ms | 2 |
| $\tau_s$ | Synapse time constant | ms | 2 |

The synaptic input *I*^syn^(*t*) from other neurons *j* to neuron *i* is modelled with an alpha function [93]:

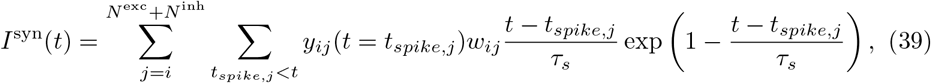

where *τ*_*s*_ is the time constant for the synapse, *t*_*spike,j*_ is the presynaptic spike time from neuron *j*, and *N* ^exc^ is the number of excitatory neurons and *N* ^inh^ is the number of inhibitory neurons. Note that all spike times include a synaptic time delay *t*_*delay*_, although this is not explicitly written. *y*_*ij*_(*t* = *t*_*spike,j*_) is the efficacy of the excitatory spike due to presynaptic short-term dynamics given by Eq (3) at the time of *t*_*spike,j*_, and *w*_*ij*_ is the postsynaptic strength that is modified by long-term plasticity. Also note that for both the SL-STDP and uncoupled TM-Triplet networks, *y*_*ij*_(*t*) is identical but *w*_*ij*_ is updated in a different manner. For inhibitory connections *y*_*ij*_(*t*) = 1 and *w*_*ij*_ is always fixed.

### Input encoding

The external input *I*^ext^(*t*) is defined in a similar manner as the synaptic input Eq (39). However, there are no short-term dynamics, i.e. *y*_*ij*_ = 1, and the spike times are generated by Poisson processes. There are two Poisson generators that result in two different external currents:

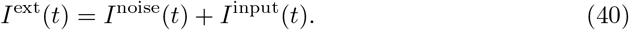

*I*^noise^(*t*) represents the static noise that is generated by a homogeneous Poisson process, and *I*^input^(*t*) represents the information capacity task related input signal that is generated by an inhomogeneous Poisson process. Each excitatory and inhibitory neuron receives these Poisson processes independently. *I*^input^(*t*) is set to zero during self-organization periods (except, see S2) and activated during the information capacity tasks.

The rates of the Poisson processes define the strength of the external current. The rates are calculated relative to a threshold rate *λ*_*θ*_, which is enough to excite an isolated neuron to emit a spike [32]:

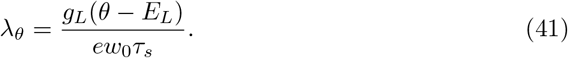

Here *e* denotes Napier’s constant and *w*_0_ is the initial input current amplitude that is calculated to be enough to cause a postsynaptic potential of strength 0.1 mV. We use a unitless proportionality multiplier *η* to define the actual input strength.

For all the self-organization tasks, we use the rate *λ*_noise_ = *η*_noise_*λ*_*θ*_ with *η*_noise_ = 3.0.

During the information capacity tasks, *η*_noise_ is lowered to 1.0 and 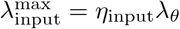 with *η*_input_ = 1.0. The input rate for the information capacity tasks is changing in time in a stepwise manner with a step size of 50 ms. The rates are uniformly distributed in the interval 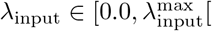.

### Network initialization

Network simulations were performed using NEST 3.8 [94] with the help of NESTML [95]. All simulations were run with a step size of 0.1 ms using 8 CPU cores.

All studied networks consists of *N*^exc^ = 800 and *N*^inh^ = 200 neurons, connected recurrently without autapses with a probability of *p* = 0.3. Synaptic time delays are randomized from a uniform probability distribution *t*_*delay*_ *∈* [0.1, 1.0[ms. Recurrent excitatory synaptic weights are initialized from a Gaussian distribution with a mean *µ*^exc^ = *w*_0_*/U*_*b*_ and standard deviation *σ*^exc^ = 0.1*µ*^exc^. Similarly, inhibitory connections are initialized from a Gaussian distribution with a mean *µ*^inh^ = − *gw*_0_ and standard deviation *σ*^inh^ = 0.1 | *µ*^inh^|. Here, *g* = 5 is the relative strength between inhibitory and excitatory synapses [32].

The noise input *I*^noise^(*t*) is connected to each neuron with a weight *w*_0_. In contrast, the input *I*^input^(*t*) is connected in a distributed manner [45]. In more detail, the input is applied to only half of the excitatory and inhibitory neurons with connection weights randomized from *w*_*ij*_ *∈* [0.0, 2*w*_0_[.

### Network analysis

The mean firing rates for each excitatory neuron were calculated after self-organization over a time window of 10 seconds. The in- and out-degree analyses were performed using NetworkX [96]. Network bursting frequencies were evaluated by first smoothing the spiking activity with a Gaussian kernel and then calculating the mean network activity. Network bursting frequency then responds to the first peak in the mean network activity’s autocorrelation function. The spike-contrast metric [62] was calculated using Elephant [97].

The mean firing rates were clustered with the k-means algorithm into five clusters in Fig 10. For the connectivity data shown in Fig 10B, Fig 10D and Fig 10E, connections with small *w* values were filtered out. Only connections with *w* at least 5% of the initial weight were preserved.

### Working memory task

The working memory tasks involve two distinct types of learning. First, the RNNs use STDP rules (SL-STDP or Triplet) to self-organize for 30 minutes (except 10 minutes for the weight normalization condition). After this, long-term plasticity is turned off and the self-organized RNNs are used for reservoir computing. The reservoir computing phase is split into two phases, a training phase and a testing phase. In the training phase, the network is simulated for *T* = 30 000 input steps, and the activity of the excitatory neurons is used to fit a linear readout. In the testing phase, the network is simulated for a new set of *T* ^*′*^ = 15 000 input steps. The predicted output signal is then calculated by using the fitted linear readout and the excitatory neurons’ activity during the test phase. Splitting the reservoir computing phase into two phases was necessary to avoid overfitting. This entire procedure is repeated 10 times with different random number seeds to reduce trial-to-trial variability in the capacity calculation.

#### Capacity computation

The instantaneous firing rate *λ*_*i*_(*t*) for each excitatory neuron *i* is approximated by convolving the neural spiking data with a Gaussian kernel of width 20 ms. The output of the system is then a linear transformation

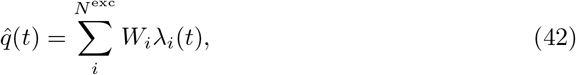

where *W*_*i*_ are the weights of the output layer. During the training phase, this estimator is fitted to a target function *q*(*t*) by minimizing the mean squared error (MSE) [42]:

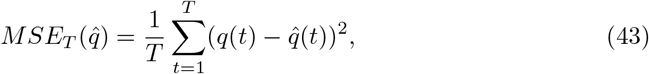

where *T* is the duration over the whole input. The training capacity of the system *C*^train^ is then calculated as [42]

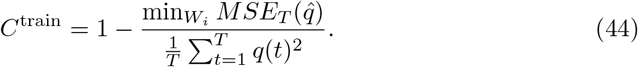

The testing capacity is calculated in a similar manner, except that the already fitted weights *W*_*i*_ are used for the computation.

#### Target generation

The output target *q*(*t*) is constructed from the input signal *λ*_input_(*t*) as a product of normalized Legendre polynomials 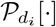 of degree *d*_*i*_ at each time step [42]:

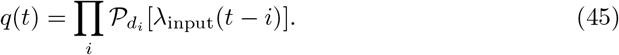

The index *i* represents the fading memory component of the transformation. As subtly described in [45], the sum of all degrees *d*_*i*_ defines the total polynomial degree and the maximum delay is the largest delay *i* for which *d*_*i*_ > 0.

As an example, the output target signal for the cell 2-0 in Fig 11 (total degree = 2 and maximum delay = 0) can be defined only in one way with a tuple (2) corresponding to an output target *q*(*t*) = *P*_2_[*λ*_input_(*t*)]. In contrast, the cell 2-1 may be defined in two different ways: a tuple (1, 1) leading to an output target signal

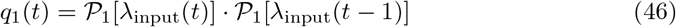

or a tuple (0, 2) leading to another output target

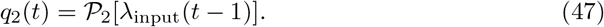

In both cases, the total degree equals 2 and the maximum delay is 1. Because of this ambiguity, we repeat the target generation for each cell for a total of 10 times favoring different combinations as much as possible. The total capacity for each cell *k* is then calculated as a weighted sum

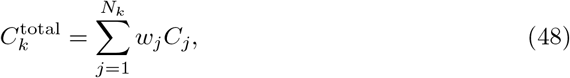

where the number of combinations *N*_*k*_ depends on the cell *k*. The weight *w*_*j*_ for each target *q*_*j*_(*t*) is decided in a round-robin fashion so that Σ_*j*_ *w*_*j*_ = 10. Consequently, the maximum total capacity for each cell in the capacity matrix is 10, since the capacity *C*_*j*_ for each *q*_*j*_(*t*) is bounded to [0, 1].

## Supporting information

Supplementary Results 6

Supplementary Results 5

Supplementary Results 4

Supplementary Results 3

Supplementary Results 2

Supplementary Results 1

## Data availability statement

All data and code used for running experiments, model fitting, and plotting is available on a GitHub repository at https://github.com/IiroAhokainen/SL-STDP.

## Supporting information

**S1 S1: Supplementary Triplet model results**

**S2 S2: Effects of initial mean firing rates on network clustering**

**S3 S3: Analysis of inhibitory activity**

**S4 S4: Supplementary information capacity results**

**S5 S5: SL-STDP model without postsynaptic gain control**

**S6 S6: Synapse models’ parameter comparison and network simulation parameters**

## Acknowledgments

We would like to thank Dr. Mikko Lehtimäki and Saana Seppälä for their insightful feedback and Dr. Charl Linssen for help with the NESTML implementation.

## References

1. Markram H, Lübke J, Frotscher M, Sakmann B. Regulation of Synaptic Efficacy by Coincidence of Postsynaptic APs and EPSPs. Science. 1997;275(5297):213–215. doi:10.1126/science.275.5297.213.

2. Bi Gq, Poo Mm. Synaptic Modifications in Cultured Hippocampal Neurons: Dependence on Spike Timing, Synaptic Strength, and Postsynaptic Cell Type. The Journal of Neuroscience. 1998;18(24):10464–10472. doi:10.1523/jneurosci.18-24-10464.1998.

3. Vogels TP, Sprekeler H, Zenke F, Clopath C, Gerstner W. Inhibitory Plasticity Balances Excitation and Inhibition in Sensory Pathways and Memory Networks. Science. 2011;334(6062):1569–1573. doi:10.1126/science.1211095.

4. Zenke F, Agnes EJ, Gerstner W. Diverse synaptic plasticity mechanisms orchestrated to form and retrieve memories in spiking neural networks. Nature Communications. 2015;6(1). doi:10.1038/ncomms7922.

5. Hiratani N, Fukai T. Interplay between Short- and Long-Term Plasticity in Cell-Assembly Formation. PLoS ONE. 2014;9(7):e101535. doi:10.1371/journal.pone.0101535.

6. Bi Gq, Poo Mm. Synaptic Modification by Correlated Activity: Hebb’s Postulate Revisited. Annual Review of Neuroscience. 2001;24(1):139–166. doi:10.1146/annurev.neuro.24.1.139.

7. Gütig R, Aharonov R, Rotter S, Sompolinsky H. Learning Input Correlations through Nonlinear Temporally Asymmetric Hebbian Plasticity. The Journal of Neuroscience. 2003;23(9):3697–3714. doi:10.1523/jneurosci.23-09-03697.2003.

8. Pfister JP, Gerstner W. Triplets of Spikes in a Model of Spike Timing-Dependent Plasticity. The Journal of Neuroscience. 2006;26(38):9673–9682. doi:10.1523/jneurosci.1425-06.2006.

9. Clopath C, Büsing L, Vasilaki E, Gerstner W. Connectivity reflects coding: a model of voltage-based STDP with homeostasis. Nature Neuroscience. 2010;13(3):344–352. doi:10.1038/nn.2479.

10. Nevian T, Sakmann B. Spine Ca2+Signaling in Spike-Timing-Dependent Plasticity. The Journal of Neuroscience. 2006;26(43):11001–11013. doi:10.1523/jneurosci.1749-06.2006.

11. Graupner M. Mechanisms of induction and maintenance of spike-timing dependent plasticity in biophysical synapse models. Frontiers in Computational Neuroscience. 2010;4. doi:10.3389/fncom.2010.00136.

12. Echeveste R, Gros C. Two-Trace Model for Spike-Timing-Dependent Synaptic Plasticity. Neural Computation. 2015;27(3):672–698. doi:10.1162/neco_a_00707.

13. Froemke RC, Dan Y. Spike-timing-dependent synaptic modification induced by natural spike trains. Nature. 2002;416(6879):433–438. doi:10.1038/416433a.

14. Froemke RC, Tsay IA, Raad M, Long JD, Dan Y. Contribution of Individual Spikes in Burst-Induced Long-Term Synaptic Modification. Journal of Neurophysiology. 2006;95(3):1620–1629. doi:10.1152/jn.00910.2005.

15. Froemke RC. Temporal modulation of spike-timing-dependent plasticity. Frontiers in Synaptic Neuroscience. 2010; doi:10.3389/fnsyn.2010.00019.

16. Shah NT, Yeung LC, Cooper LN, Cai Y, Shouval HZ. A Biophysical Basis for the Inter-spike Interaction of Spike-timing-dependent Plasticity. Biological Cybernetics. 2006;95(2):113–121. doi:10.1007/s00422-006-0071-y.

17. Schmiedt J, Albers C, Pawelzik K. Spike timing-dependent plasticity as dynamic filter. In: Lafferty J, Williams C, Shawe-Taylor J, Zemel R, Culotta A, editors. Advances in Neural Information Processing Systems. vol. 23. Curran Associates, Inc.; 2010.Available from: https://proceedings.neurips.cc/paper_files/paper/2010/file/892c91e0a653ba19df81a90f89d99bcd-Paper.pdf.

18. Albers C, Schmiedt JT, Pawelzik KR. Theta-specific susceptibility in a model of adaptive synaptic plasticity. Frontiers in Computational Neuroscience. 2013;7. doi:10.3389/fncom.2013.00170.

19. Deperrois N, Graupner M. Short-term depression and long-term plasticity together tune sensitive range of synaptic plasticity. PLOS Computational Biology. 2020;16(9):e1008265. doi:10.1371/journal.pcbi.1008265.

20. Cai Y, Gavornik JP, Cooper LN, Yeung LC, Shouval HZ. Effect of Stochastic Synaptic and Dendritic Dynamics on Synaptic Plasticity in Visual Cortex and Hippocampus. Journal of Neurophysiology. 2007;97(1):375–386. doi:10.1152/jn.00895.2006.

21. Kumar A, Mehta MR. Frequency-Dependent Changes in NMDAR-Dependent Synaptic Plasticity. Frontiers in Computational Neuroscience. 2011;5. doi:10.3389/fncom.2011.00038.

22. Chindemi G, Abdellah M, Amsalem O, Benavides-Piccione R, Delattre V, Doron M, et al. A calcium-based plasticity model for predicting long-term potentiation and depression in the neocortex. Nature Communications. 2022;13(1). doi:10.1038/s41467-022-30214-w.

23. Ecker A, Egas Santander D, Abdellah M, Alonso JB, Bolaños-Puchet S, Chindemi G, et al. Assemblies, synapse clustering, and network topology interact with plasticity to explain structure-function relationships of the cortical connectome. eLife. 2025;13. doi:10.7554/elife.101850.3.

24. Costa RP, Froemke RC, Sjöström PJ, van Rossum MC. Unified pre- and postsynaptic long-term plasticity enables reliable and flexible learning. eLife. 2015;4. doi:10.7554/elife.09457.

25. Barak O, Tsodyks M. Working models of working memory. Current Opinion in Neurobiology. 2014;25:20–24. doi:10.1016/j.conb.2013.10.008.

26. Wang XJ. 50 years of mnemonic persistent activity: quo vadis? Trends in Neurosciences. 2021;44(11):888–902. doi:10.1016/j.tins.2021.09.001.

27. Mongillo G, Barak O, Tsodyks M. Synaptic Theory of Working Memory. Science. 2008;319(5869):1543–1546. doi:10.1126/science.1150769.

28. Kozachkov L, Tauber J, Lundqvist M, Brincat SL, Slotine JJ, Miller EK. Robust and brain-like working memory through short-term synaptic plasticity. PLOS Computational Biology. 2022;18(12):e1010776. doi:10.1371/journal.pcbi.1010776.

29. Fitz H, Uhlmann M, van den Broek D, Duarte R, Hagoort P, Petersson KM. Neuronal spike-rate adaptation supports working memory in language processing. Proceedings of the National Academy of Sciences. 2020;117(34):20881–20889. doi:10.1073/pnas.2000222117.

30. Tsodyks M, Pawelzik K, Markram H. Neural Networks with Dynamic Synapses. Neural Computation. 1998;10(4):821–835. doi:10.1162/089976698300017502.

31. Sjöström PJ, Turrigiano GG, Nelson SB. Rate, Timing, and Cooperativity Jointly Determine Cortical Synaptic Plasticity. Neuron. 2001;32(6):1149–1164. doi:10.1016/s0896-6273(01)00542-6.

32. Brunel N. Dynamics of Sparsely Connected Networks of Excitatory and Inhibitory Spiking Neurons. Journal of Computational Neuroscience. 2000;8(3):183–208. doi:10.1023/a:1008925309027.

33. Watt AJ. Homeostatic plasticity and STDP: keeping a neuron’s cool in a fluctuating world. Frontiers in Synaptic Neuroscience. 2010;2. doi:10.3389/fnsyn.2010.00005.

34. Oja E. Simplified neuron model as a principal component analyzer. Journal of Mathematical Biology. 1982;15(3):267–273. doi:10.1007/bf00275687.

35. Goodhill GJ, Barrow HG. The Role of Weight Normalization in Competitive Learning. Neural Computation. 1994;6(2):255–269. doi:10.1162/neco.1994.6.2.255.

36. Turrigiano GG. The Self-Tuning Neuron: Synaptic Scaling of Excitatory Synapses. Cell. 2008;135(3):422–435. doi:10.1016/j.cell.2008.10.008.

37. Lu Jt, Li Cy, Zhao JP, Poo Mm, Zhang Xh. Spike-Timing-Dependent Plasticity of Neocortical Excitatory Synapses on Inhibitory Interneurons Depends on Target Cell Type. The Journal of Neuroscience. 2007;27(36):9711–9720. doi:10.1523/jneurosci.2513-07.2007.

38. Kullmann DM, Lamsa KP. LTP and LTD in cortical GABAergic interneurons: Emerging rules and roles. Neuropharmacology. 2011;60(5):712–719. doi:10.1016/j.neuropharm.2010.12.020.

39. Kuhlman SJ, Olivas ND, Tring E, Ikrar T, Xu X, Trachtenberg JT. A disinhibitory microcircuit initiates critical-period plasticity in the visual cortex. Nature. 2013;501(7468):543–546. doi:10.1038/nature12485.

40. Bannon NM, Chistiakova M, Volgushev M. Synaptic Plasticity in Cortical Inhibitory Neurons: What Mechanisms May Help to Balance Synaptic Weight Changes? Frontiers in Cellular Neuroscience. 2020;14. doi:10.3389/fncel.2020.00204.

41. Grier BD, Parkins S, Omar J, Lee HK. Selective plasticity of fast and slow excitatory synapses on somatostatin interneurons in adult visual cortex. Nature Communications. 2023;14(1). doi:10.1038/s41467-023-42968-y.

42. Dambre J, Verstraeten D, Schrauwen B, Massar S. Information Processing Capacity of Dynamical Systems. Scientific Reports. 2012;2(1). doi:10.1038/srep00514.

43. Maass W, Natschläger T, Markram H. Real-Time Computing Without Stable States: A New Framework for Neural Computation Based on Perturbations. Neural Computation. 2002;14(11):2531–2560. doi:10.1162/089976602760407955.

44. Verstraeten D, Schrauwen B, D’Haene M, Stroobandt D. An experimental unification of reservoir computing methods. Neural Networks. 2007;20(3):391–403. doi:10.1016/j.neunet.2007.04.003.

45. Schulte to Brinke T, Dick M, Duarte R, Morrison A. A refined information processing capacity metric allows an in-depth analysis of memory and nonlinearity trade-offs in neurocomputational systems. Scientific Reports. 2023;13(1). doi:10.1038/s41598-023-37604-0.

46. Tsodyks M, Markram H. The neural code between neocortical pyramidal neurons depends on neurotransmitter releaseprobability. Proceedings of the National Academy of Sciences. 1997;94(2):719–723. doi:10.1073/pnas.94.2.719.

47. Markram H, Wang Y, Tsodyks M. Differential signaling via the same axon of neocortical pyramidal neurons. Proceedings of the National Academy of Sciences. 1998;95(9):5323–5328. doi:10.1073/pnas.95.9.5323.

48. Costa RP, Sjostrom PJ, van Rossum MC. Probabilistic inference of short-term synaptic plasticity in neocortical microcircuits. Frontiers in Computational Neuroscience. 2013;Volume 7 – 2013. doi:10.3389/fncom.2013.00075.

49. Ghanbari A, Malyshev A, Volgushev M, Stevenson IH. Estimating short-term synaptic plasticity from pre- and postsynaptic spiking. PLOS Computational Biology. 2017;13(9):e1005738. doi:10.1371/journal.pcbi.1005738.

50. Johnston D, Wu SMs, Gray R. Foundations of cellular neurophysiology. Cambridge, Mass: MIT Press; 1995.

51. Budde T, Meuth S, Pape HC. Calcium-dependent inactivation of neuronal calcium channels. Nature Reviews Neuroscience. 2002;3(11):873–883. doi:10.1038/nrn959.

52. Limpitikul WB, Dick IE. Inactivation of CaV1 and CaV2 channels. Journal of General Physiology. 2025;157(2). doi:10.1085/jgp.202313531.

53. Magee JC, Johnston D. A Synaptically Controlled, Associative Signal for Hebbian Plasticity in Hippocampal Neurons. Science. 1997;275(5297):209–213. doi:10.1126/science.275.5297.209.

54. Johnston D, Christie BR, Frick A, Gray R, Hoffman DA, Schexnayder LK, et al. Active dendrites, potassium channels and synaptic plasticity. Philosophical Transactions of the Royal Society of London Series B: Biological Sciences. 2003;358(1432):667–674. doi:10.1098/rstb.2002.1248.

55. Bienenstock E, Cooper L, Munro P. Theory for the development of neuron selectivity: orientation specificity and binocular interaction in visual cortex. The Journal of Neuroscience. 1982;2(1):32–48. doi:10.1523/jneurosci.02-01-00032.1982.

56. Senn W, Markram H, Tsodyks M. An Algorithm for Modifying Neurotransmitter Release Probability Based on Pre- and Postsynaptic Spike Timing. Neural Computation. 2001;13(1):35–67. doi:10.1162/089976601300014628.

57. Thomson AM, Deuchars J, West DC. Large, deep layer pyramid-pyramid single axon EPSPs in slices of rat motor cortex display paired pulse and frequency-dependent depression, mediated presynaptically and self-facilitation, mediated postsynaptically. Journal of Neurophysiology. 1993;70(6):2354–2369. doi:10.1152/jn.1993.70.6.2354.

58. Thomson AM. Activity-dependent properties of synaptic transmission at two classes of connections made by rat neocortical pyramidal axons in vitro. The Journal of Physiology. 1997;502(1):131–147. doi:10.1111/j.1469-7793.1997.131bl.x.

59. Litwin-Kumar A, Doiron B. Formation and maintenance of neuronal assemblies through synaptic plasticity. Nature Communications. 2014;5(1). doi:10.1038/ncomms6319.

60. Graupner M, Brunel N. Calcium-based plasticity model explains sensitivity of synaptic changes to spike pattern, rate, and dendritic location. Proceedings of the National Academy of Sciences. 2012;109(10):3991–3996. doi:10.1073/pnas.1109359109.

61. Fardet T, Ballandras M, Bottani S, Métens S, Monceau P. Understanding the Generation of Network Bursts by Adaptive Oscillatory Neurons. Frontiers in Neuroscience. 2018;12. doi:10.3389/fnins.2018.00041.

62. Ciba M, Isomura T, Jimbo Y, Bahmer A, Thielemann C. Spike-contrast: A novel time scale independent and multivariate measure of spike train synchrony. Journal of Neuroscience Methods. 2018;293:136–143. doi:10.1016/j.jneumeth.2017.09.008.

63. Holt GR, Softky WR, Koch C, Douglas RJ. Comparison of discharge variability in vitro and in vivo in cat visual cortex neurons. Journal of Neurophysiology. 1996;75(5):1806–1814. doi:10.1152/jn.1996.75.5.1806.

64. Vinck M, Batista-Brito R, Knoblich U, Cardin J. Arousal and Locomotion Make Distinct Contributions to Cortical Activity Patterns and Visual Encoding. Neuron. 2015;86(3):740–754. doi:10.1016/j.neuron.2015.03.028.

65. Hoy JL, Niell CM. Layer-Specific Refinement of Visual Cortex Function after Eye Opening in the Awake Mouse. The Journal of Neuroscience. 2015;35(8):3370–3383. doi:10.1523/jneurosci.3174-14.2015.

66. Verstraeten D, Dambre J, Dutoit X, Schrauwen B. Memory versus non-linearity in reservoirs. In: The 2010 International Joint Conference on Neural Networks (IJCNN). IEEE; 2010. p. 1–8. Available from: 10.1109/IJCNN.2010.5596492.

67. Reyes A, Lujan R, Rozov A, Burnashev N, Somogyi P, Sakmann B. Target-cell-specific facilitation and depression in neocortical circuits. Nature Neuroscience. 1998;1(4):279–285. doi:10.1038/1092.

68. Angulo MC, Staiger JF, Rossier J, Audinat E. Distinct Local Circuits Between Neocortical Pyramidal Cells and Fast-Spiking Interneurons in Young Adult Rats. Journal of Neurophysiology. 2003;89(2):943–953. doi:10.1152/jn.00750.2002.

69. Silberberg G, Markram H. Disynaptic Inhibition between Neocortical Pyramidal Cells Mediated by Martinotti Cells. Neuron. 2007;53(5):735–746. doi:10.1016/j.neuron.2007.02.012.

70. Wang Y, Markram H, Goodman PH, Berger TK, Ma J, Goldman-Rakic PS. Heterogeneity in the pyramidal network of the medial prefrontal cortex. Nature Neuroscience. 2006;9(4):534–542. doi:10.1038/nn1670.

71. Kiritani T, Wickersham IR, Seung HS, Shepherd GMG. Hierarchical Connectivity and Connection-Specific Dynamics in the Corticospinal–Corticostriatal Microcircuit in Mouse Motor Cortex. The Journal of Neuroscience. 2012;32(14):4992–5001. doi:10.1523/jneurosci.4759-11.2012.

72. Kawaguchi Y. Pyramidal Cell Subtypes and Their Synaptic Connections in Layer 5 of Rat Frontal Cortex. Cerebral Cortex. 2017;27(12):5755–5771. doi:10.1093/cercor/bhx252.

73. Martens MB, Houweling AR, E Tiesinga PH. Anti-correlations in the degree distribution increase stimulus detection performance in noisy spiking neural networks. Journal of Computational Neuroscience. 2016;42(1):87–106. doi:10.1007/s10827-016-0629-1.

74. Winding M, Pedigo BD, Barnes CL, Patsolic HG, Park Y, Kazimiers T, et al. The connectome of an insect brain. Science. 2023;379(6636). doi:10.1126/science.add9330.

75. Bae JA, Baptiste M, Baptiste MR, Bishop CA, Bodor AL, Brittain D, et al. Functional connectomics spanning multiple areas of mouse visual cortex. Nature. 2025;640(8058):435–447. doi:10.1038/s41586-025-08790-w.

76. Vasilaki E, Giugliano M. Emergence of Connectivity Motifs in Networks of Model Neurons with Short- and Long-Term Plastic Synapses. PLOS ONE. 2014;9(1):e84626. doi:10.1371/journal.pone.0084626.

77. Abeles M. Corticonics: Neural Circuits of the Cerebral Cortex. Cambridge University Press; 1991. Available from: 10.1017/CBO9780511574566.

78. Lazar A. SORN: a Self-organizing Recurrent Neural Network. Frontiers in Computational Neuroscience. 2009;3. doi:10.3389/neuro.10.023.2009.

79. Zenke F, Gerstner W. Hebbian plasticity requires compensatory processes on multiple timescales. Philosophical Transactions of the Royal Society B: Biological Sciences. 2017;372(1715):20160259. doi:10.1098/rstb.2016.0259.

80. Hennequin G, Agnes EJ, Vogels TP. Inhibitory Plasticity: Balance, Control, and Codependence. Annual Review of Neuroscience. 2017;40(1):557–579. doi:10.1146/annurev-neuro-072116-031005.

81. Mackwood O, Naumann LB, Sprekeler H. Learning excitatory-inhibitory neuronal assemblies in recurrent networks. eLife. 2021;10. doi:10.7554/elife.59715.

82. van Rossum MCW, Bi GQ, Turrigiano GG. Stable Hebbian Learning from Spike Timing-Dependent Plasticity. The Journal of Neuroscience. 2000;20(23):8812–8821. doi:10.1523/jneurosci.20-23-08812.2000.

83. Gilson M, Fukai T. Stability versus Neuronal Specialization for STDP: Long-Tail Weight Distributions Solve the Dilemma. PLoS ONE. 2011;6(10):e25339. doi:10.1371/journal.pone.0025339.

84. Buzsáki G, Mizuseki K. The log-dynamic brain: how skewed distributions affect network operations. Nature Reviews Neuroscience. 2014;15(4):264–278. doi:10.1038/nrn3687.

85. Yu Q, Tsodyks M, Sompolinsky H, Schmitz D, Gütig R. Interactions between long- and short-term synaptic plasticity transform temporal neural representations into spatial. Proceedings of the National Academy of Sciences. 2025;122(47). doi:10.1073/pnas.2426290122.

86. Gütig R, Sompolinsky H. The tempotron: a neuron that learns spike timing–based decisions. Nature Neuroscience. 2006;9(3):420–428. doi:10.1038/nn1643.

87. Kudithipudi D, Schuman C, Vineyard CM, Pandit T, Merkel C, Kubendran R, et al. Neuromorphic computing at scale. Nature. 2025;637(8047):801–812. doi:10.1038/s41586-024-08253-8.

88. Gjorgjieva J, Clopath C, Audet J, Pfister JP. A triplet spike-timing–dependent plasticity model generalizes the Bienenstock–Cooper–Munro rule to higher-order spatiotemporal correlations. Proceedings of the National Academy of Sciences. 2011;108(48):19383–19388. doi:10.1073/pnas.1105933108.

89. Byrd RH, Lu P, Nocedal J, Zhu C. A Limited Memory Algorithm for Bound Constrained Optimization. SIAM Journal on Scientific Computing. 1995;16(5):1190–1208. doi:10.1137/0916069.

90. Zhu C, Byrd RH, Lu P, Nocedal J. Algorithm 778: L-BFGS-B: Fortran subroutines for large-scale bound-constrained optimization. ACM Transactions on Mathematical Software. 1997;23(4):550–560. doi:10.1145/279232.279236.

91. Wales DJ, Doye JPK. Global Optimization by Basin-Hopping and the Lowest Energy Structures of Lennard-Jones Clusters Containing up to 110 Atoms. The Journal of Physical Chemistry A. 1997;101(28):5111–5116. doi:10.1021/jp970984n.

92. Meurer A, Smith CP, Paprocki M, Čertík O, Kirpichev SB, Rocklin M, et al. SymPy: symbolic computing in Python. PeerJ Computer Science. 2017;3:e103. doi:10.7717/peerj-cs.103.

93. De Schutter E. Computational modeling methods for neuroscientists. Computational neuroscience. MIT Press; 2010.

94. Graber S, Mitchell J, Kurth AC, Terhorst D, Skaar JEW, Schöfmann CM, et al. NEST 3.8; 2024. Available from: 10.5281/zenodo.12624784.

95. Linssen C, Babu PN, Rumpe B, Morrison A. NESTML 8.0.1; 2025. Available from: 10.5281/zenodo.15292719.

96. Hagberg AA, Schult DA, Swart PJ. Exploring Network Structure, Dynamics, and Function using NetworkX. In: Varoquaux G, Vaught T, Millman J, editors. Proceedings of the 7th Python in Science Conference. Pasadena, CA USA; 2008. p. 11–15.

97. Denker M, Yegenoglu A, Grün S. Collaborative HPC-enabled workflows on the HBP Collaboratory using the Elephant framework. In: Neuroinformatics 2018; 2018. p. P19. Available from: https://abstracts.g-node.org/conference/NI2018/abstracts#/uuid/023bec4e-0c35-4563-81ce-2c6fac282abd.

