## Supplementary Results 6 for "A unified model of short- and long-term plasticity: Effects on network connectivity and information capacity"

### S6: Synapse model parameters

Iiro Ahokainen and Marja-Leena Linne, Tampere University

**Table 1.** Comparison of depressing synapse parameter values for the SL-STDP model and the uncoupled TM-Triplet model network simulations. These parameters are used in all E-E connections and also depressing E $\rightarrow$ I connections. In simulations where E $\rightarrow$ I connections are not plastic, then $f = 0$.

| Parameter | Description | SL-STDP | Uncoupled TM-Triplet | Units |
|---|---|---|---|---|
| $\tau_r$ | Depression time constant | 200 | 200 | ms |
| $\tau_f$ | Facilitation time constant | 50 | 50 | ms |
| $U$ | Release probability increment per presynaptic spike | 0.2029 | 0.2029 | – |
| $U_b$ | Baseline release probability | 0.7019 | 0.7019 | – |
| $\tau_y$ | Time constant of presynaptic spike efficacy $y(t)$ | 16.54 | 16.54 | ms |
| $\tau_c$ | Time constant of Triplet presynaptic trace $c(t)$ | – | 16.8 | ms |
| $\tau_-$ | Time constant of LTD postsynaptic trace $z_1(t)$ | 100.0 | 33.7 | ms |
| $\tau_+$ | Time constant of LTP postsynaptic trace $z_2(t)$ | 63.30 | 35.09 | ms |
| $Z_1$ | Gain control of LTD postsynaptic trace | 0.4102 | – | – |
| $Z_2$ | Gain control of LTP postsynaptic trace | 0.1158 | – | – |
| $A_-$ | LTD update amplitude | 0.0166 | 0.0076 | – |
| $A_+$ | LTP update amplitude | 0.8499 | 0.0285 | – |
| $f$ | Learning rate | 10.96 | 10.96 | pA |

**Table 2.** Comparison of facilitating synapse parameter values for the SL-STDP model and the uncoupled TM-Triplet model network simulations. These parameters are used in facilitating E $\rightarrow$ I connections.

| Parameter | Description | SL-STDP | Uncoupled TM-Triplet | Units |
|---|---|---|---|---|
| $\tau_r$ | Depression time constant | 50 | 50 | ms |
| $\tau_f$ | Facilitation time constant | 500 | 500 | ms |
| $U$ | Release probability increment per presynaptic spike | 0.15 | 0.15 | – |
| $U_b$ | Baseline release probability | 0.15 | 0.15 | – |
| $\tau_y$ | Time constant of presynaptic spike efficacy $y(t)$ | 16.54 | 16.54 | ms |
| $\tau_c$ | Time constant of Triplet presynaptic trace $c(t)$ | – | 16.8 | ms |
| $\tau_-$ | Time constant of LTD postsynaptic trace $z_1(t)$ | 100.0 | 33.7 | ms |
| $\tau_+$ | Time constant of LTP postsynaptic trace $z_2(t)$ | 63.30 | 35.09 | ms |
| $Z_1$ | Gain control of LTD postsynaptic trace | 0.4102 | – | – |
| $Z_2$ | Gain control of LTP postsynaptic trace | 0.1158 | – | – |
| $A_-$ | LTD update amplitude | 0.0166 | 0.0076 | – |
| $A_+$ | LTP update amplitude | 0.8499 | 0.0285 | – |
| $f$ | Learning rate | 10.96 | 10.96 | pA |

### Network simulation parameters

**Table 3.** Network simulation specific parameters

| Parameter | Description | Value | Units |
|---|---|---|---|
| $w_{\text{init}}$ | Initial postsynaptic strength | $10.96 \pm 1.096^{\dagger}$ | pA |
| $w_{\text{max}}$ | Maximal postsynaptic strength | 109.6 | pA |
| $t_{\text{delay}}$ | Spike time delay | [0.1, 1.0[* | ms |
| $p$ | Initial connection probability | 0.3 | - |
| $\lambda_{\theta}$ | Threshold Poisson noise rate | 8968 | Hz |

† Sampled from Gaussian distribution mean±std.
* Sampled from uniform distribution.
