## Supplementary Results 5 for "A unified model of short- and long-term plasticity: Effects on network connectivity and information capacity"

### S5: SL-STDP model without postsynaptic gain control

Iiro Ahokainen and Marja-Leena Linne, Tampere University

In the original Triplet model, the presynaptic and postsynaptic traces could increase without bounds. It is not obvious which biological processes could (or even should) be linked to the synaptic traces, but unlimited growth seems implausible for any biological pathway. Motivated by the limitations of resources, we included gain control also for the postsynaptic side for the SL-STDP model. This raises the question of whether the improved performance in the working memory tasks presented in the main article is due to sharing the short-term presynaptic state with long-term plasticity or to postsynaptic gain control.

To assess the contribution of sharing the latent state to the increased information capacity, we repeated the model fitting procedure and the working memory classification for the SL-STDP model without postsynpatic gain control. In other words, the model equations now read as follows:

$$\frac{dx(t)}{dt} = \frac{1-x(t)}{\tau_r} - x(t)u(t)R_{pre}(t) \tag{1}$$

$$\frac{du(t)}{dt} = \frac{U_b - u(t)}{\tau_f} + U(1-u(t))R_{pre}(t) \tag{2}$$

$$\frac{dy(t)}{dt} = -\frac{y(t)}{\tau_y} + x(t)u(t)R_{pre}(t) \tag{3}$$

$$\frac{dz_1(t)}{dt} = -\frac{z_1(t)}{\tau_-} + R_{post}(t) \tag{4}$$

$$\frac{dz_2(t)}{dt} = -\frac{z_2(t)}{\tau_+} + R_{post}(t). \tag{5}$$

The $w$ update equations are unchanged.

We follow the same fitting procedure as introduced in Methods section. We perform the model fitting for two sets of short-term dynamics parameters. First, using the same parameters as before: $\tau_f = 50$ ms and $\tau_r = 200$ ms, and secondly, slightly less depression: $\tau_f = 50$ ms and $\tau_r = 100$ ms. The fitted parameters are shown in Table 1.

**Table 1. Fitted parameters for the SL-STDP model without postsynaptic gain.**

| $\tau_r$ | $U$ | $U_b$ | $\tau_y$ (ms) | $\tau_-$ (ms) | $A^-$ | $\tau_+$ (ms) | $A^+$ | Error |
|---|---|---|---|---|---|---|---|---|
| 200 | 0.2718 | 0.8700 | 19.97 | 100.0 | 0.0058 | 62.09 | 0.0870 | 526.1 |
| 100 | 0.010 | 0.900 | 17.01 | 100.0 | 0.0053 | 45.70 | 0.0909 | 543.0 |

The short-term dynamics time constants were fixed during fitting: $\tau_r = 200$ ms (first row) or $\tau_r = 100$ ms (second row) and $\tau_f = 50$ ms for both parameter sets. All other parameters were fitted. The last column on the right shows the total fitting error.

After fitting, we continue to plot the mean drift of $w$, which now simply reads as

$$\begin{aligned}\left\langle \frac{dw}{dt} \right\rangle &= -A^-\lambda_x < z_1 > +A^+\lambda_z < z_2 >< y > \\ &= -A^-\lambda_x\tau_-\lambda_z + A^+\tau_y\tau_+\lambda_z^2 \frac{\lambda_x^2 + \frac{U_b}{\tau_f U}\lambda_x}{\frac{1}{\tau_f U} + \lambda_x(1+\frac{\tau_r U_b}{\tau_f U}) + \tau_r\lambda_x^2}.\end{aligned} \tag{6}$$

Approximating $\lambda_x << \frac{U_b}{\tau_f U}$, we find that $\left\langle \frac{dw}{dt} \right\rangle = 0$ approach to a linear dependency $\lambda_z \propto a\lambda_x + b$, where the slope $a = \frac{A^- \tau_- \tau_r}{A^+ \tau_y \tau_+}$ and the constant term $b = \frac{A^- \tau_-}{A^+ \tau_y \tau_+ U_b}$. Eq (6) with Table 1 values is illustrated in Fig 1. The line $\lambda_z = a\lambda_x + b$ defines the balance between the LTP and LTD zones. A higher slope $a$ favors LTD-dominant dynamics. For a strongly facilitating synapse, the previous approximation may not hold.

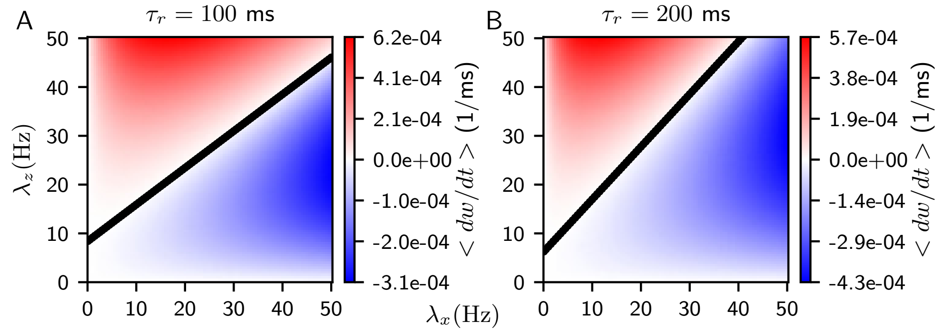


**Fig 1. The mean drift $\left\langle \frac{dw}{dt} \right\rangle$ for the SL-STDP model without gain control.** Eq (6) is plotted as a function of presynaptic firing rate $\lambda_x$ and postsynaptic firing rate $\lambda_z$. (**A**) $\tau_r = 100$ ms and (**B**) $\tau_r = 200$ ms, with parameter values from Table 1. The red area shows the net LTP regime, and the blue area shows the net LTD regime. The white area shows balanced regions where $\left\langle \frac{dw}{dt} \right\rangle \approx 0$. The black line is the line separating LTP and LTD zones $\lambda_z = a\lambda_x + b$.

Similar to the SL-STDP model with postsynaptic gain control, we observe that there are two local maxima (or "optimal update zones") that depend non-trivially on $\lambda_x$. With $\tau_r = 200$ ms (Fig 1B), the LTD zone dominates over the LTP zone, whereas $\tau_r = 100$ ms (Fig 1A) shows a more balanced distribution between the zones. Indeed, repeating the self-organization study presented in the main article leads to weakening of the majority of synapses in the case $\tau_r = 200$ ms, as shown in Fig 2C-D. In contrast, the case $\tau_r = 100$ ms produces clustering similar to the postsynaptic gain controlled SL-STDP model (Fig 2A-B).

In particular, there is one more cluster in the case $\tau_r = 100$ ms than is seen for the model with the postsynaptic gain (Fig 6A-B and Fig 10 in the main article). Furthermore, the maximal mean firing rates are elevated. However, the correlation trends for both out-degrees and in-degrees remain unchanged.

Next, we repeated the working memory task for both cases, $\tau_r = 200$ ms and $\tau_r = 100$ ms. The results are shown in Fig 3. Surprisingly, both systems performed well, although the in- and out-degree distributions are markedly different. The LTD-dominant system (Fig 3B) shows particularly good performance in simple input transformations (cells 1-2 and 2-1 responding to a single input time step delay and a quadratic transformation without time delay, respectively). However, for more complex transformations, i.e. time delay of four time steps and a quartic transformation, the balanced LTP-LTD system shows better performance (Fig 3A).

Taken together, these results indicate that, for RNNs, the exact form of the synaptic update equations is not the dominant factor. Instead, it is the emerging balance between the LTP and LTD zones that is critical. The SL-STDP model, with or without postsynaptic gain control, is able to produce different dynamic balances between the LTP and LTD regions, and this balance correlates with the information capacity of the

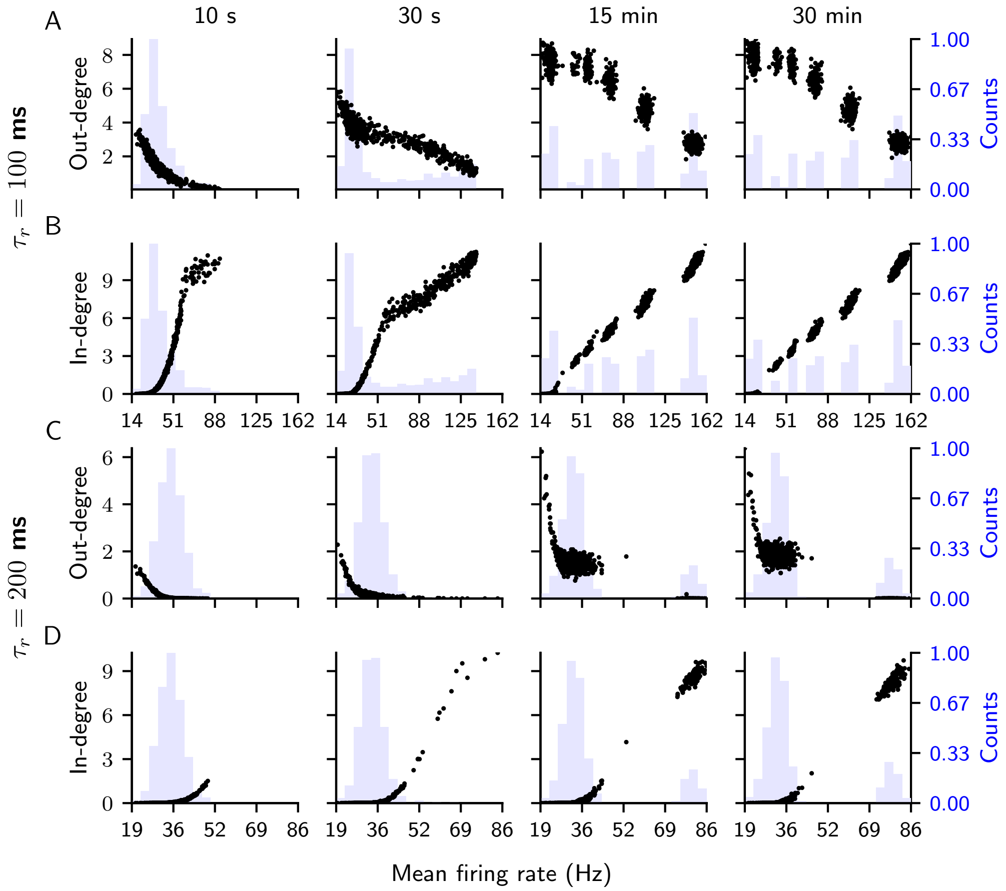


**Fig 2. Evolution of relative total out-degree and in-degree strengths with respect to mean firing rates for the SL-STDP model without postsynaptic gain control.** Panels from left to right show the evolution of weight strengths from 10 seconds to 30 minutes of self-organization with respect to a mean firing rate of each excitatory neuron. Histograms of the mean firing rates are shown in light blue. (**A**, **C**) Total out-degree strength relative to the initial total out-degree for each of the $N_e = 800$ excitatory neurons in the SL-STDP network, shown as black dots. Each normalized total out-degree is plotted against the mean firing rate of the same neuron. (**B**, **D**) Similar as (**A**, **C**), but for the relative total in-degree strength. In (**A-B**), parameters of the second row of Table 1 are used, while (**C-D**) uses the parameters of the first row.

network.

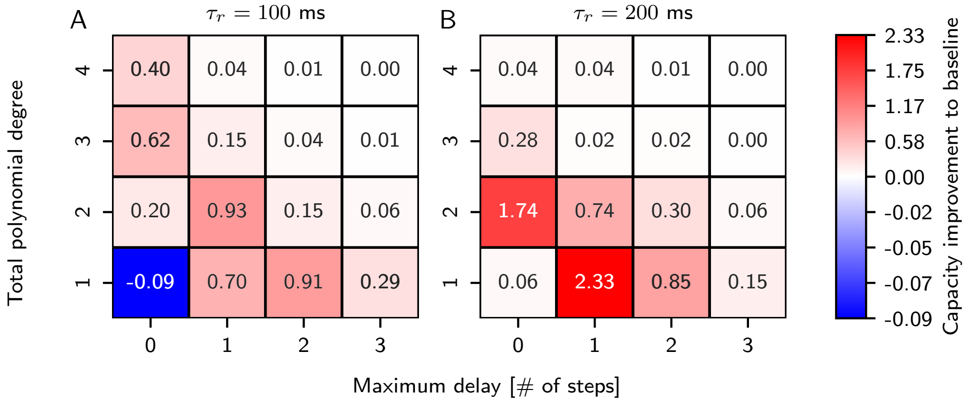


**Fig 3. Working memory capacities for the SL-STDP model without postsynaptic gain control.** (**A**) Model with balanced LTP/LTD regions ($\tau_r = 100$ ms). (**B**) Model with LTD-dominant parameters ($\tau_r = 200$ ms). Higher values indicate better output prediction. Increasing total polynomial degree and maximum delay corresponds to more difficult working memory tasks. The reported values are mean testing capacities averaged over ten different random initializations of the networks.
