## Supplementary Results 4 for "A unified model of short- and long-term plasticity: Effects on network connectivity and information capacity"

### S4: Supplementary information capacity results

Iiro Ahokainen and Marja-Leena Linne, Tampere University

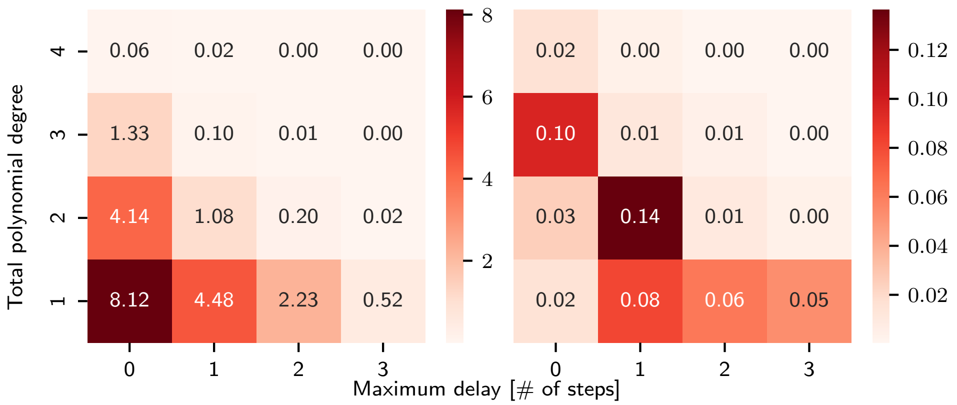


**Fig 1. Absolute test capacity values for the baseline network.** (**Left**) Mean capacity values over 10 different random initializations of the network. A capacity value of 10 is the theoretical maximum. Increasing total polynomial degree and maximum delay correspond to more difficult working memory tasks. (**Right**) Corresponding standard deviations for each cell shown on the left.

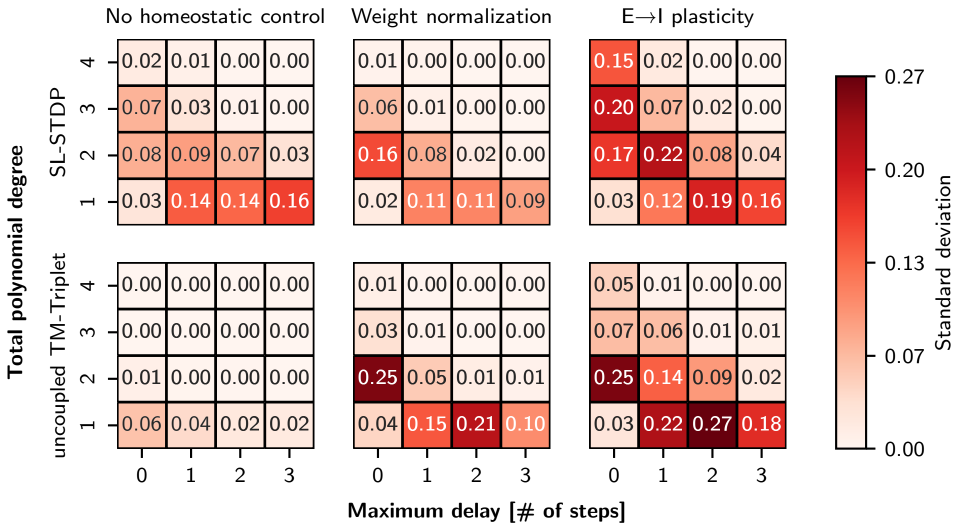


**Fig 2. Standard deviations of test capacities with different homeostatic mechanisms.** The standard deviations of the absolute capacity values over 10 runs are shown for each combination of plasticity model and homeostatic control mechanism are showed.

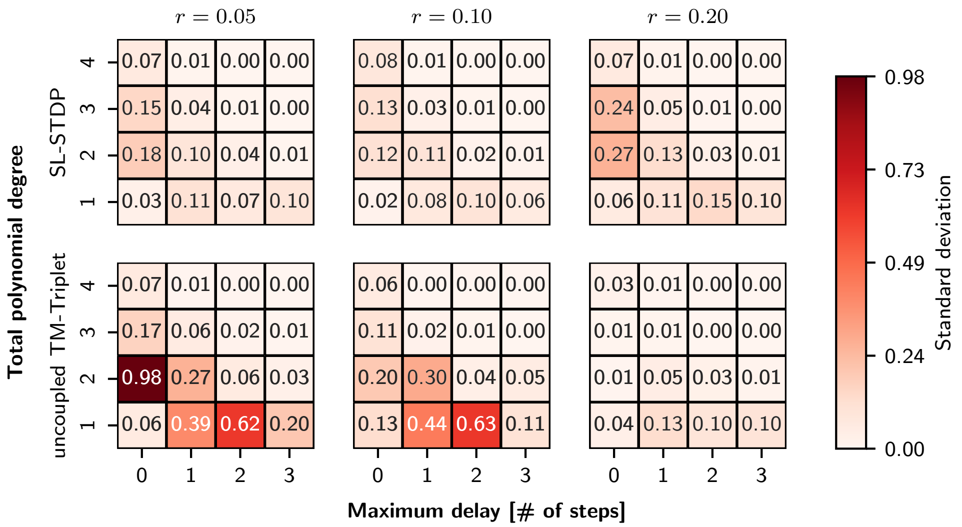


**Fig 3. Standard deviations of test capacities with different ratios of facilitating and depressing synapses.** The standard deviations of the absolute capacity values over 10 runs are shown for each combination of plasticity model and ratio $r$ between facilitating and depressing $\mathrm{E} \rightarrow \mathrm{I}$ synapses. Overall, the SL-STDP model shows less deviation than the uncoupled TM-Triplet model in almost all cells.
