## Supplementary Results 3 for "A unified model of short- and long-term plasticity: Effects on network connectivity and information capacity"

### S3: Analysis of inhibitory activity

Iiro Ahokainen and Marja-Leena Linne, Tampere University

This supplementary section presents additional analyses of inhibitory network activity for both the SL-STDP and uncoupled TM-Triplet models under different self-organization conditions. We characterize inhibitory dynamics using firing rate statistics and coefficient-of-variation (CV) distributions. The results highlight differences in inhibitory firing regularity and self-organization between the two plasticity models, particularly in the presence of excitatory-to-inhibitory plasticity.

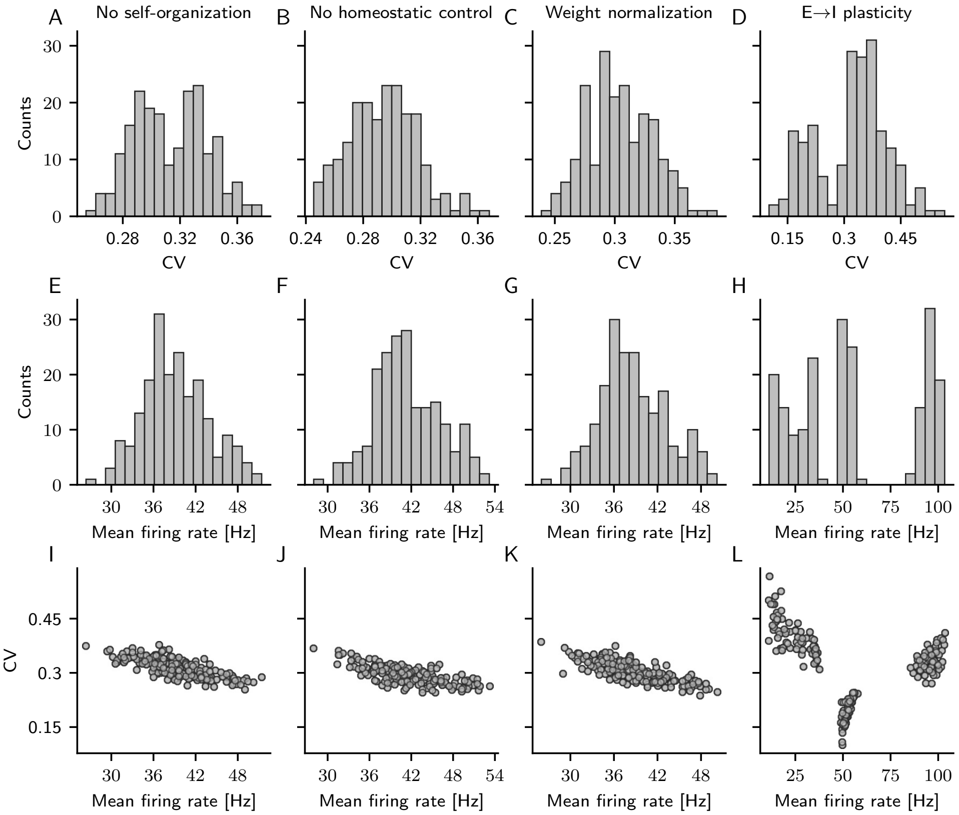


**Fig 1. SL-STDP networks' inhibitory activity statistics with different conditions.** The top row (**A-D**) shows Coefficient of Variation (CV) histograms for all inhibitory neurons evaluated over five seconds of network activity. The middle row (**E-H**) shows the mean firing rates for all inhibitory neurons. The bottom row (**I-L**) shows the CV values as a function of the mean firing rates. The first column shows the network statistics prior to self-organization, meaning that the same statistics also apply to the uncoupled TM-Triplet model before self-organization. The mean firing rates of inhibitory neurons remain fairly similar when no homeostatic control or weight normalization is applied (**E-G**). (**H**) In contrast, E $\rightarrow$ I plasticity clusters the firing rates mimicking excitatory neurons' activity (see Fig 9 in the main article), and the CV histogram shows two peaks (**D**). (**L**) Each cluster has preferred range of CV values. Interestingly, the lowest CV range is with the cluster around 50 Hz. Note that this coincides with the network oscillation frequency.

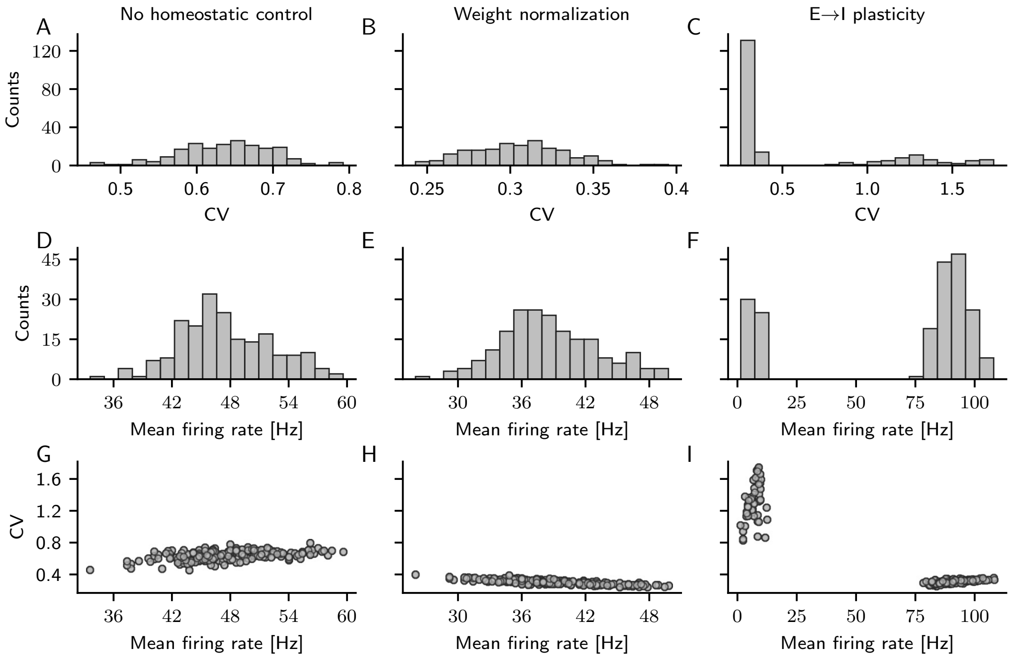


**Fig 2. Uncoupled TM-Triplet networks' inhibitory activity statistics with different conditions.** The top row (**A-C**) shows Coefficient of Variation (CV) histograms for all inhibitory neurons evaluated over five seconds of network activity. The middle row (**D-F**) shows the mean firing rates for all inhibitory neuron. The bottom row (**G-I**) shows the CV values as a function of the mean firing rates. CV values for the case without homeostatic control (**A**) are elevated compared to the SL-STDP network (Fig 1B). (**F**) E $\rightarrow$ I plasticity clusters inhibitory firing rates into two clusters following the excitatory neurons' activity (see Fig 9 in the main article). (**C**) Small amount of CV values reach over one, indicating irregularity in the firing.
