## Supplementary Results 2 for "A unified model of short- and long-term plasticity: Effects on network connectivity and information capacity"

### S2: Effects of initial mean firing rates on network clustering

Iiro Ahokainen and Marja-Leena Linne, Tampere University

In the main article, we performed the in- and out-degree analysis using relatively high input noise ($\eta = 3.0$), resulting in mean firing rates of $\lambda_{\mathrm{exc}} \approx \lambda_{\mathrm{inh}} = 39$ Hz. Such high firing rates rarely occur for prolonged periods in the cortex, especially in excitatory neurons [1–3]. This raises the question of whether the same dynamics emerge at lower baseline mean firing rates. To address this, we performed a series of tests to see how the network self-organizes under lower mean firing rates.

To test whether the network can still self-organize into distinct firing-rate groups, as shown in Fig 6A–B of the main article, at lower mean firing rates, we repeated the same self-organization analysis with the input noise reduced by half ($\eta = 1.5$). In this case, the initial mean firing rates are $\lambda_{\mathrm{exc}} \approx \lambda_{\mathrm{inh}} = 14$ Hz, i.e. the network is net depressing, as predicted by the mean-field analysis in Fig 4B in the main article. Indeed, simulations with $\eta = 1.5$ reveal that there is insufficient net potentiation for clustering to occur in this input firing rate regime. Instead, all connections decouple from each other after approximately 1 minute of simulation.

The difficulty arises from the use of homogeneous Poisson processes for the stimulation, which leads to steady firing rates across all neurons. If we instead repeat the self-organization anlysis by applying temporally changing inputs, we get more realistic situation. To test this, we repeated the self-organization analysis using the same input settings as in the working memory task (see the Methods section in the main article). With this setup, $\eta = 1.5$ results in $\lambda_{\mathrm{exc}} \approx 21$ Hz and $\lambda_{\mathrm{inh}} \approx 22$ Hz, while $\eta = 1.2$ yields $\lambda_{\mathrm{exc}} \approx 14$ Hz and $\lambda_{\mathrm{inh}} \approx 15$ Hz. The resulting in- and out-degree distributions are shown in Fig 1.

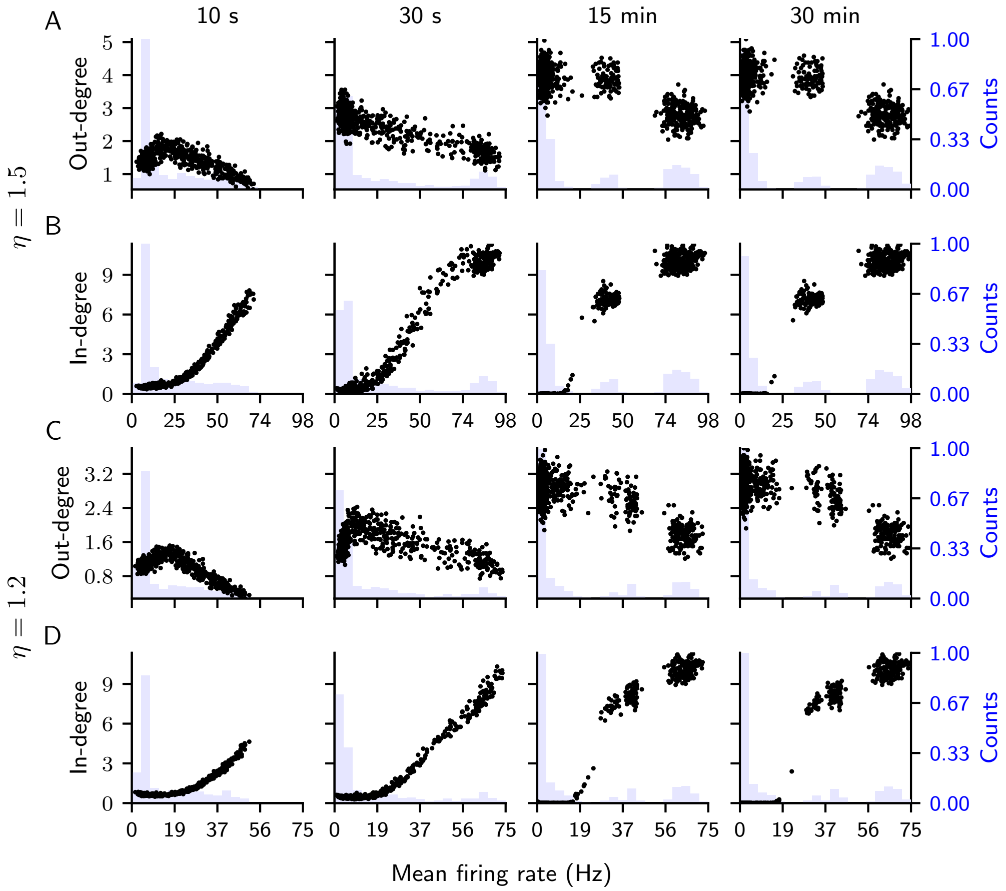


**Fig 1. Evolution of relative total out-degree and in-degree strengths with respect to mean firing rates.** Panels from left to right show the evolution of weight strengths from 10 seconds to 20 minutes of self-organization with respect to a mean firing rate of each excitatory neuron. Histograms of the mean firing rates are shown in light blue. (**A**, **C**) Total out-degree strength relative to initial total out-degree for each of $N_e = 800$ excitatory neurons in the SL-STDP network, shown as black dots. Each normalized total out-degree is plotted against the mean firing rate of the same neuron. (**B**, **D**) Similar as (**A**, **C**) but for the relative total in-degree strength. In (**A-B**), the input strength $\eta$ is set to 1.5 while in (**C-D**) $\eta = 1.2$.

Fig 1 shows that clustering is still present at lower baseline firing rates, although the fraction of nodes in the middle clusters decreases as the average input strength is reduced. We do not have a descriptive theory for this dependence of the number and size of the clusters on the initial baseline mean firing rates, as the appearance of the middle clusters were not predicted in the first place. We are not aware of any model or theory that would explain this interaction, and thus leave a more detailed analysis for future studies.

Similarly to the main article, we calculate the maximal synchrony $S$ values for both cases. After self-organization, we find $S = 0.15$ with bin size $\Delta = 14.5$ ms for $\eta = 1.5$, and $S = 0.09$ with bin size $\Delta = 15.9$ ms for $\eta = 1.2$. The initial coefficient of variation of the excitatory population changed from 0.64 to 0.69 after self-organization for the $\eta = 1.5$ case. A raster plot and the spike-contrast metric [4] after 30 minutes of self-organization are shown in Fig 2 for the case $\eta = 1.5$. From Fig 2A we see, and indicated by the non-zero synchrony value, that the network oscillations still exist for the stepwise changing inputs. However, the oscillations are less synchronous than before.

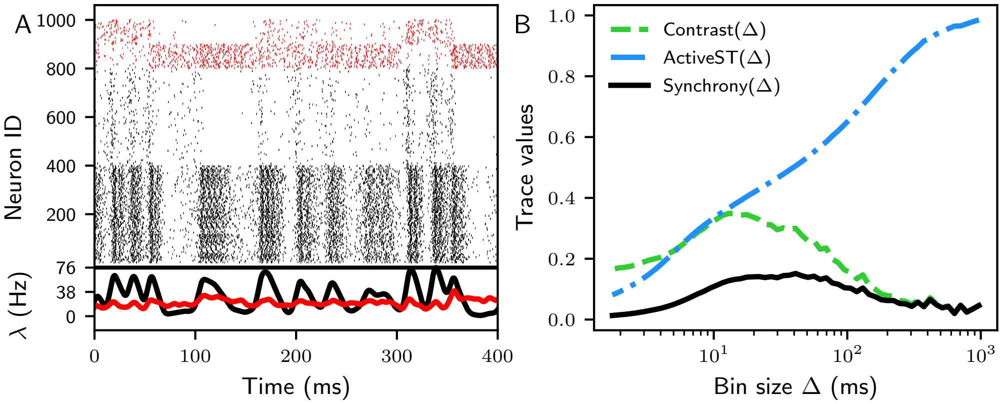


**Fig 2. Activity of the SL-STDP network after 30 minutes of self-organization, exhibiting bursting and synchronization.** (**A**) Raster plot of the SL-STDP network after 30 minutes of self-organization. Black markers indicate spike times of excitatory neurons, and red markers indicate spike times of inhibitory neurons. The first 400 excitatory neurons and inhibitory neurons 800-899 receive alternating inputs. At the bottom, the averaged population activity $\lambda$ is shown for excitatory neurons (black) and inhibitory neurons (red). The excitatory population exhibits oscillations, while the inhibitory population remains largely asynchronous. (**B**) Spike-contrast metric [4] for the excitatory spikes shown in (**A**). Synchrony is measured as a product of Contrast and ActiveST (active spike trains). The maximum synchrony of 0.15 is achieved with a bin size $\Delta = 14.5$ ms.

#### References

1. Holt GR, Softky WR, Koch C, Douglas RJ. Comparison of discharge variability in vitro and in vivo in cat visual cortex neurons. Journal of Neurophysiology. 1996;75(5):1806–1814. doi:10.1152/jn.1996.75.5.1806.

2. Vinck M, Batista-Brito R, Knoblich U, Cardin J. Arousal and Locomotion Make Distinct Contributions to Cortical Activity Patterns and Visual Encoding. Neuron. 2015;86(3):740–754. doi:10.1016/j.neuron.2015.03.028.

3. Hoy JL, Niell CM. Layer-Specific Refinement of Visual Cortex Function after Eye Opening in the Awake Mouse. The Journal of Neuroscience. 2015;35(8):3370–3383. doi:10.1523/jneurosci.3174-14.2015.

4. Ciba M, Isomura T, Jimbo Y, Bahmer A, Thielemann C. Spike-contrast: A novel time scale independent and multivariate measure of spike train synchrony. Journal of Neuroscience Methods. 2018;293:136–143. doi:10.1016/j.jneumeth.2017.09.008.
