## Supplementary Results 1 for "A unified model of short- and long-term plasticity: Effects on network connectivity and information capacity"

### S1: Supplementary Triplet model results

Iiro Ahokainen and Marja-Leena Linne, Tampere University

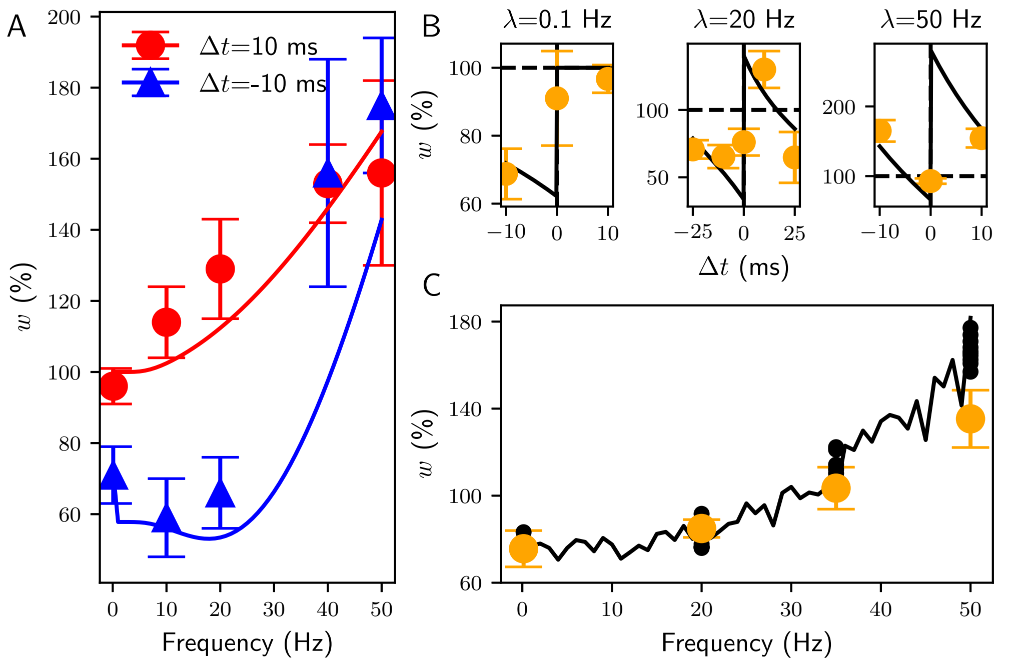


**Fig 1. Minimal all-to-all triplet model [1] fitted to the long-term plasticity induction protocol** [2]. As in [1], $\tau_c = 16.8$ ms and $\tau_- = 33.7$ ms were fixed, and the remaining parameters were fitted. The fitted parameters and the fitting error $E$ are: $A^+ = 0.0285$, $A^- = 0.0076$, $\tau_+ = 35.09$ ms, and $E = 914.8$. (**A**) Change in $w$ in response to varying interspike intervals, resulting in a nonlinear frequency response with fixed $\Delta t = \pm 10$. Dots and triangles, with standard deviations, represent data from [2], and the lines indicate the fits. (**B**) The update window with varying $\Delta t$ at three different frequencies (0.1 Hz, 20 Hz, and 50 Hz), shown from left to right. (**C**) Model fit to the randomized $\Delta t$. As in [2], $\Delta t$ was drawn from a Gaussian distribution with a mean of zero and a standard deviation of 7 ms. Orange error dots show the experimental data, and black dots show all the realizations of model fits. The black line shows one example fit across the entire frequency range.

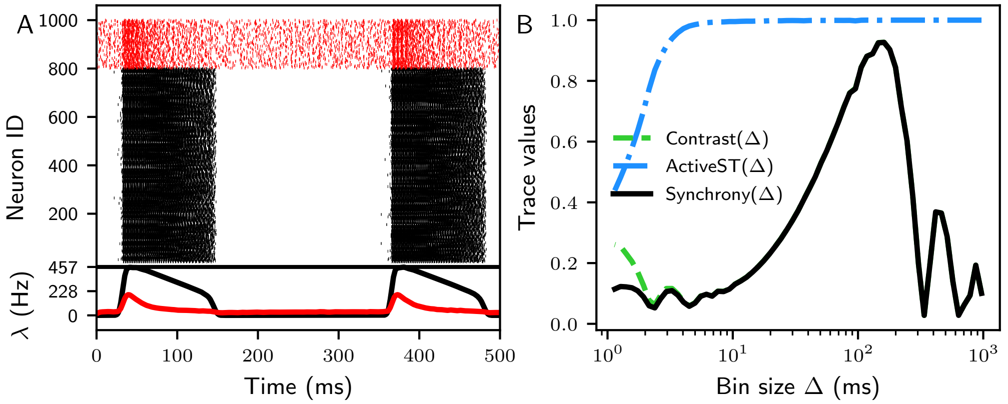


**Fig 2. Activity of the uncoupled TM-Triplet network after 30 minutes of self-organization, exhibiting network bursting and synchronization.** (**A**) Raster plot of the uncoupled TM-Triplet network after 30 minutes of self-organization. Black markers indicate spike times of excitatory neurons, and red markers indicate spike times of inhibitory neurons. At the bottom, the averaged population activity $\lambda$ is shown for excitatory neurons (black) and inhibitory neurons (red). Due to the runaway potentiation of E-E connections, the uncoupled TM-Triplet network exhibits superficially high firing rates for long times during bursts. Both the excitatory and inhibitory populations show bursting activity. (**B**) Spike-contrast metric [3] for the excitatory spikes shown in (**A**). Synchrony is measured as a product of Contrast and ActiveST (active spike trains). The maximum synchrony of 0.93 is achieved with a bin size of $\Delta = 145.4$ ms.

### References

1. Pfister JP, Gerstner W. Triplets of Spikes in a Model of Spike Timing-Dependent Plasticity. The Journal of Neuroscience. 2006;26(38):9673–9682. doi:10.1523/jneurosci.1425-06.2006.

2. Sjöström PJ, Turrigiano GG, Nelson SB. Rate, Timing, and Cooperativity Jointly Determine Cortical Synaptic Plasticity. Neuron. 2001;32(6):1149–1164. doi:10.1016/s0896-6273(01)00542-6.

3. Ciba M, Isomura T, Jimbo Y, Bahmer A, Thielemann C. Spike-contrast: A novel time scale independent and multivariate measure of spike train synchrony. Journal of Neuroscience Methods. 2018;293:136–143. doi:10.1016/j.jneumeth.2017.09.008.
